# STABILITY ANALYSIS OF A SIGNALING CIRCUIT WITH DUAL SPECIES OF GTPASE SWITCHES

**DOI:** 10.1101/2020.08.31.276311

**Authors:** Lucas M. Stolerman, Pradipta Ghosh, Padmini Rangamani

## Abstract

GTPases are molecular switches that regulate a wide range of cellular processes, such as organelle biogenesis, position, shape, and function, vesicular transport between organelles, and signal transduction. These hydrolase enzymes operate by toggling between an active “ON”) guanosine triphosphate (GTP)-bound state and an inactive (“OFF”) guanosine diphosphate (GDP)-bound state; such a toggle is regulated by GEFs (guanine nucleotide exchange factors) and GAPs (GTPase activating proteins). Here we propose a model for a network motif between monomeric (m) and trimeric (t) GTPases assembled exclusively in eukaryotic cells of multicellular organisms. We develop a system of ordinary differential equations in which these two classes of GTPases are interlinked conditional to their ON/OFF states within a motif through coupling and feedback loops. We provide explicit formulae for the steady states of the system and perform classical local stability analysis to systematically investigate the role of the different connections between the GTPase switches. Interestingly, a coupling of the active mGTPase to the GEF of the tGTPase was sufficient to provide two locally stable states: one where both active/inactive forms of the mGTPase can be interpreted as having low concentrations and the other where both m- and tGTPase have high concentrations. Moreover, when a feedback loop from the GEF of the tGTPase to the GAP of the mGTPase was added to the coupled system, two other locally stable states emerged, both having the tGTPase inactivated and being interpreted as having low active tGTPase concentrations. Finally, the addition of a second feedback loop, from the active tGTPase to the GAP of the mGTPase, gives rise to a family of steady states that can be parametrized by a range of inactive tGTPase concentrations. Our findings reveal that the coupling of these two different GTPase motifs can dramatically change their steady state behaviors and shed light on how such coupling may impact signaling mechanisms in eukaryotic cells.

## 1. Introduction

Each eukaryotic cell has many a large number of GTP-binding proteins (also called *GTPases* or G-proteins). They are thought to be intermediates in an extended cellular signaling and transport network that touches on nearly every aspect of cell function [1, 2, 3]. One unique feature of GTPases is that they serve as *biochemical switches* that exist in an ‘OFF’ state when bound to a guanosine diphosphate (GDP), and can be turned ‘ON’ when that GDP is exchanged for a guanosine triphosphate (GTP) nucleotide [1, 4]. Turning the GTPase ‘ON’ is the key rate limiting step in the activation-inactivation process, requires an external stimulus, and is catalyzed by a class of enzymes called guanine nucleotide exchange factors (GEFs) [5]. G proteins return to their ‘OFF’ state when the bound GTP is hydrolyzed to guanosine diphosphate (GDP) via an intrinsic hydrolase activity of the GTPase; this step is catalyzed by GTPase-activating proteins (GAPs) [6]. Thus, GEFs and GAPs play a crucial role in controlling the dynamics of the GTPase switch and the finiteness of signaling that it transduces [7, 8, 9, 10]. Dysregulation of GTPase switches has been implicated in cellular malfunctioning and is commonly encountered in diverse diseases [11, 12, 13, 14]. For example, hyperactivation of GTPases [15, 16] is known to support a myriad of cellular phenotypes that contribute to aggressive tumor traits [17, 18]. Such traits have also been associated with aberrant activity of GAPs [15] or GEFs [19, 20, 21, 22, 23, 24]. These works underscore the importance of GTPases as vital regulators of high fidelity cellular communication.

There are two distinct types of GTPases that gate signals: small or monomeric (m) and trimeric (t) GTPases. mGTPases are mostly believed to function within the cell’s interior and are primarily concerned with organelle function and cytoskeletal remodeling [25, 26, 27]. tGTPases, on the other hand, were believed to primarily function at the cell’s surface from where they gate the duration, type and extent of signals that are initiated by receptors on the cell’s surface [28, 29]. These two classes of switches were believed to function largely independently, until early 1990’s when tGTPases were detected on intracellular membranes, e.g., the Golgi [29, 30], and studies alluded to the possibility that they, alongside mGTPases, may co-regulate organelle function and structure [31]. But it was not until 2016 that the first evidence of an example of functional coupling between the two switches – m- and tGTPases– emerged. Using a combination of biochemical, biophysical, structural modeling, live cell imaging, and numerous read-outs of Golgi functions, it was shown that m- and tGTPase co-regulate each other on the Golgi [32]. The specific discovery of GIV/Girdin, a non-receptor GEF for G*αi*, as a platform for crosstalk between trimeric G proteins and monomeric Arf1 GTPases at the Golgi is the main biological motivation of the present study. In Fig 1A, we depict where these proteins interact in the cell and what experimentally-determined sequence of events, segregated in space and time, enable the execution of key steps in secretion through the Golgi. The experiments in [32] showed that when mGTPase (Arf family) is turned ‘ON’, it engages with a GEF for tGPTase (GIV/Girdin; tGEF); the latter binds and activates tGTPase, of the Gi subfamily, G*α*i. The engaged tGEF triggers the activation of a tGTPase (Gi). Upon activation, the tGTPases activate the GAP for Arf1, ArfGAP2/3 (mGAP), via the release of ‘free’ G*βγ*. The mGAP turns ‘OFF’ the mGTPase Arf1, thereby terminating the mGTPase signaling. Termination of the mGTPase (Arf1) activity results in a finite lifetime of the Arf1 signal. This “finiteness” of signal from Arf1 is critical for membrane trafficking and organelle structure [33, 34, 35]. Thus, this phenomenon of co-regulation between the two classes of GTPases was shown to be critical in limiting the duration of mGTPase and tGTPase signaling on the Golgi membrane, which in turn significantly regulates Golgi shape and function. In doing so, this dual GTPase circuit converted simple chemical signals into complex mechanical outputs such as membrane trafficking. Emerging evidence from protein-protein interaction networks and decades of work on both species of GTPases suggest that such co-regulation through coupling between the GTPases is possible and likely occurs on multiple organellar membranes. What advantages do two coupled species of GTPase switches provide over independent, uncoupled switches? The answer to this question has not yet been experimentally dissected or intuitively theorized.

**Figure 1.**
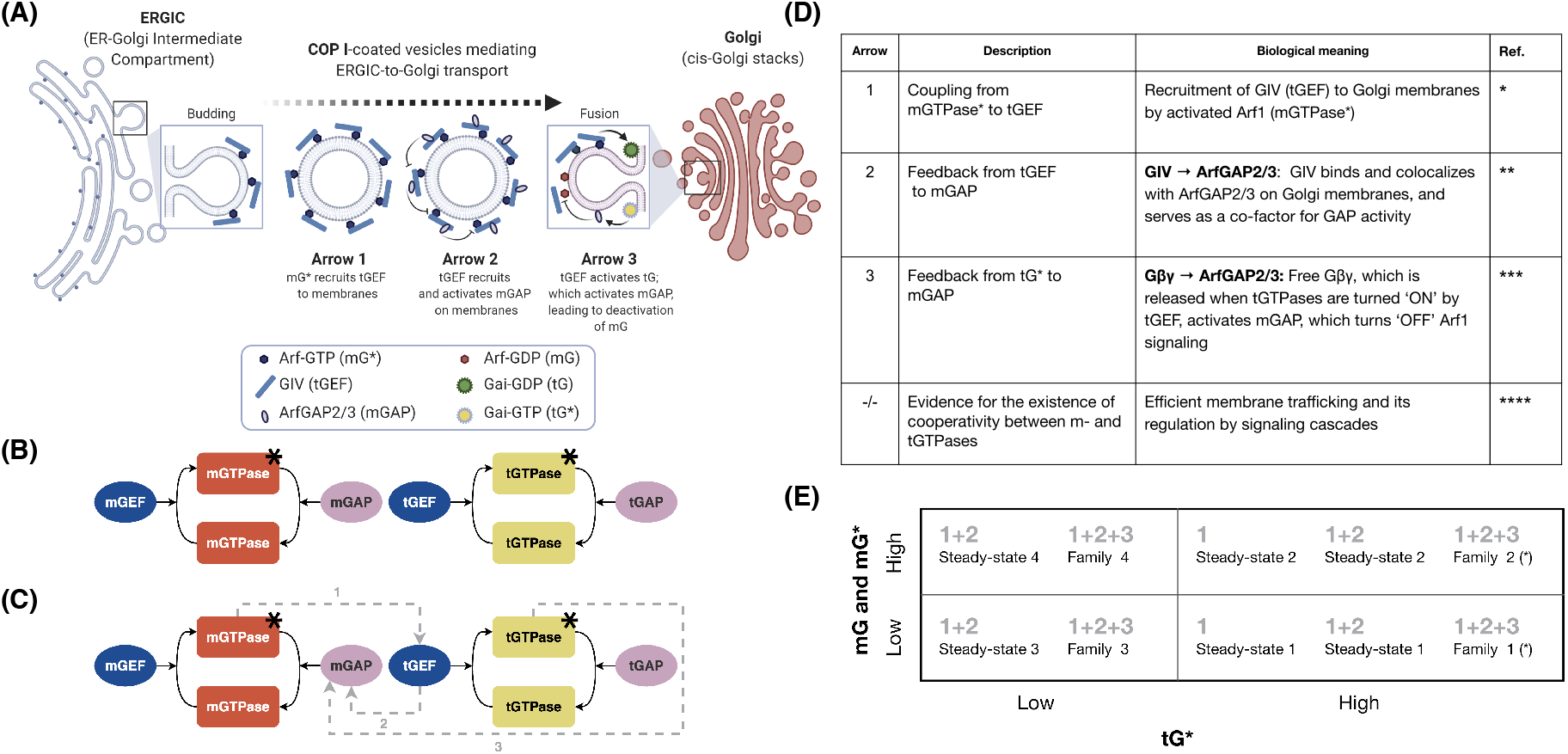
A network motif in which two species of GTPases are inter-linked. (A) Recent experimental findings revealed that monomeric Arf GTPases and trimeric G proteins co-regulate each other on the Golgi membrane (B) Uncoupled monomeric and trimeric GTPase switches are represented by mGTPase and tGTPase, respectively. The black star denotes the active forms. Activation and inactivation are regulated by GEFs and GAPs, where the first letter (m or t) indicates the associated GTPase. (C) Our proposed mathematical model describes the interaction between the two GTPase switches. Arrows 1, 2, and 3 show the coupling and feedback loops that were found experimentally. (D) Description and biological meaning of each arrow connecting the GTPase switches. References: [32] for arrow 1 (*), [32] for arrow 2 (**), [32, 63] for arrow 3 (***), and [63, 30, 64, 65, 31] for evidence of cooperativity between m and tGTPases (****). (E) For the three combinations of arrows (1, 1+2, and 1+2+3) chosen in our study, we calculate the steady state solutions for the coupled GTPase circuit model.

Mathematical models of signaling networks have contributed significantly to our understanding of how network motifs might function [36, 37, 38, 39, 40, 41]. Continuous-time dynamical systems, commonly represented by systems of ordinary differential equations (ODEs), are powerful tools for building rich and insightful mathematical models [42]. For example, a comprehensive steady state analysis of ODE system helped frame the concept of “zero-order ultrasensitivity” where large responses in the active fraction of a protein of interest are driven by small changes in the saturated ratio of the enzymes [43]. Similarly, modeling biochemical networks with dynamical systems also has revealed the existence of bistable switches and biological oscillators within a feedback network architecture [44, 45, 46]. The simple system proposed by Ferrell and Xiong [46] served as a basis for modeling cellular all-or-none responses, and hence, crucial for decision-making within several signaling processes. Dynamical systems have also be used for mapping chemical reactions into differential equations; when numerically integrated (clustered) these systems can be used to predict the evolution of large reaction networks [47, 48, 49]. For example, clustering methods have revealed the existence of recurrent structures, the so-called “network motifs’, the dynamics of which have been inferred with a Boolean kinetic system of differential equations [49]. These models produced various dynamic features corresponding to the motif structure, which allowed the understanding of underlying biology (i.e., gene expression patterns). Furthermore, studies using network ODE models have revealed that information is processed in cells through intricate connections between signaling pathways rather than individual motifs [50, 51]. From a systems biology modeling perspective, large systems of differential equations are usually hard to analyze, but when combined with experiments, they can give rise to quantitative predictions [52, 53, 54, 55, 56, 57, 58, 59, 60].

Here we built a mathematical model to investigate the stability properties of the coupled m- and tGTPase switches, the first example of its kind, that has been observed experimentally [32]. Beginning with the uncoupled GTPase switches (Fig.1B) as our starting point, we specifically sought to understand the stability features of the coupled motif (Fig.1C). We proposed a system of ODEs and obtained the steady states of this new network motif to understand the input-output relationships. Given the model formulation and the fact that we do not know the various kinetic parameters, obtaining these states is critical to our understanding of this network behavior. Then we studied the dynamic behavior under small perturbation around these steady states using local stability analysis [61, 62]. We investigated the different coupling and feedback loops between these two motifs all representing the observed biochemical and biophysical events during signal transduction (Fig.1D). In Fig.1E, we summarize the steady-states of the system. Our analyses revealed the existence of steady-states and their stability depends on the network connectivity. In particular, the coupling between the two switches through the connection between mG* and tGEF (represented by “1”) allowed for the emergence of two steady states with low/high mG and mG* concentrations, while tG* steady state concentration remained high in both cases. On the other hand, low tG* steady state concentrations were obtained when the feedback loop *tGEF* → *mGAP* was added to the system (represented by “1+2”). Finally, the feedback loop *tG*^∗^ → *mGAP* allowed for the emergence of four parametrized families of steady state within the same low/high configurations. In what follows, we present the model assumptions and derivation in §Section 2, the local stability analysis and numerical simulations in §Section 3, and discuss our findings in the context of GTPase signalling networks in §Section 4.

## 2. Model Development

In this section, we introduce our mathematical model for the GTPase coupled circuit (Fig.1C). We begin with outlining the model assumptions in §Section 2.1 and describe the reactions and governing equations in detail in §Section 2.2.

### 2.1 Assumptions

Our model describes the time evolution of the concentrations for the different system components. Table 1 contains the set of reactions in our system. We explicitly allow for three connections between the m- and the t-GTPase switches based on experimental findings [32] as described below.

**Table 1.**
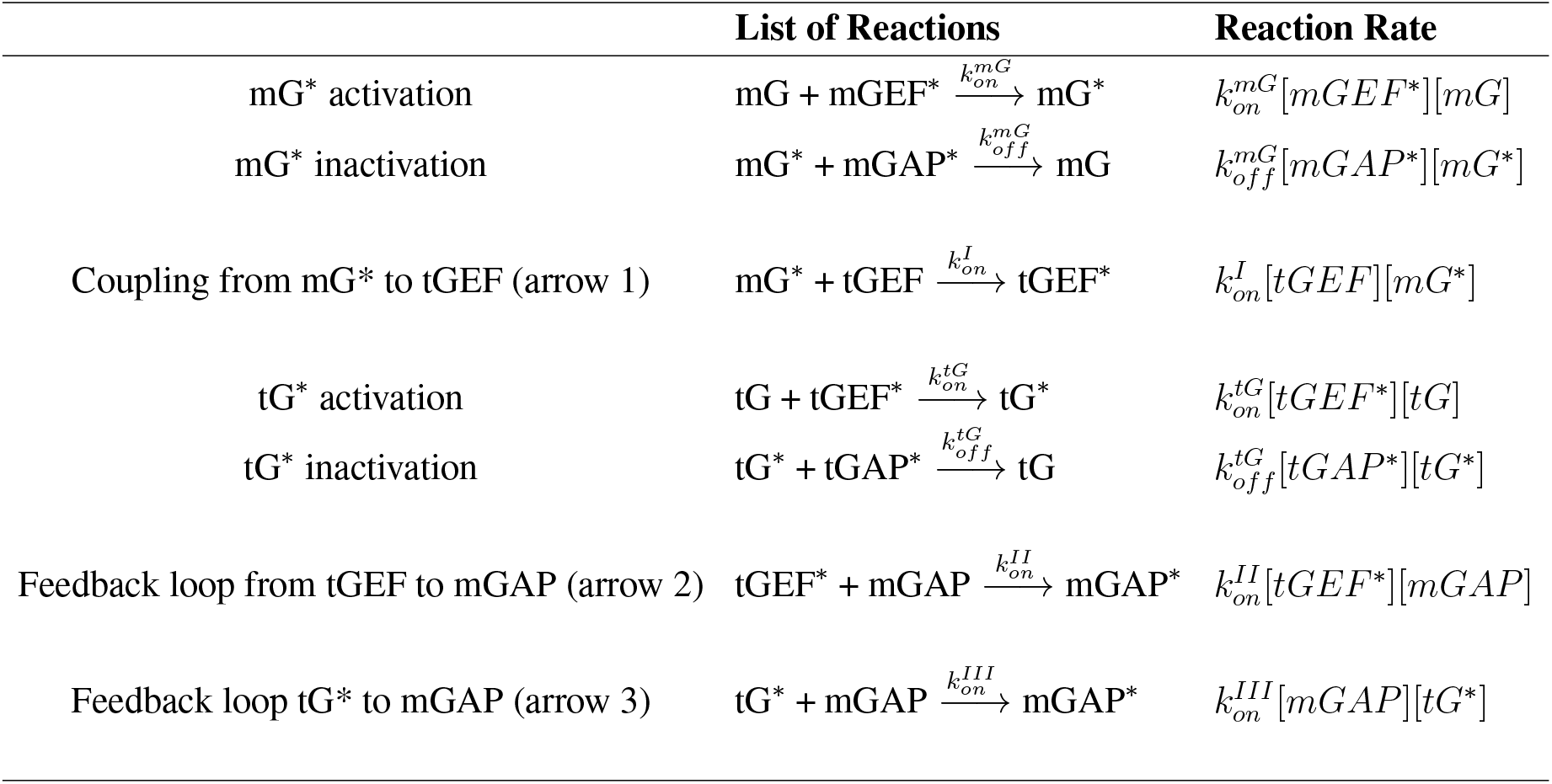
GTPase circuit reactions and rates used in the model

- Arrow 1: Represents the coupling of the two switches and represents the recruitment/engagement of tGEF by mG* (Figure 1A,B).
- Arrow 2: Represents the feedback from tGEF to mGAP (Figure 1A,B).
- Arrow 3: Represents the feedback from tG* to mGAP (Figure 1A,B).

Additionally, we only consider the toggling of GTPases that are mediated by GEFs and GAPs that activate and inactivate them, respectively.

To develop the model equations, we considered a well-mixed regime and that the concentrations of the species are in large enough amounts that deterministic kinetics hold [66, 67]. Finally, for mathematical tractability, all reactions in the system are modeled using mass-action kinetics and nonlinear kinetics such as Hill functions or Michaelis-Menten are not considered [68]. The reactions in the coupled circuit and the corresponding reaction rates used in the model are shown in Table 1.

### 2.2. Governing Equations

We developed a system of ODEs that describe coupled toggling of two switches, i.e., cyclical activation and inactivation of monomeric and trimeric GTPases within the network motif shown in Fig.1C and described in Fig.1D. In what follows, the brackets represent concentrations, which are nonnegative real numbers. The system of equations are given by

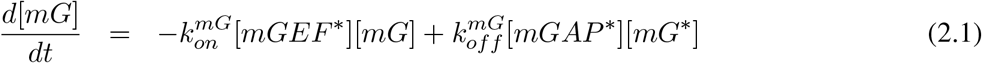

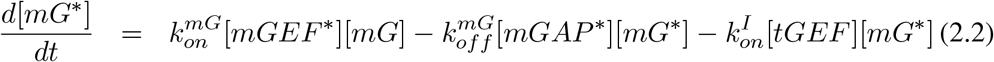

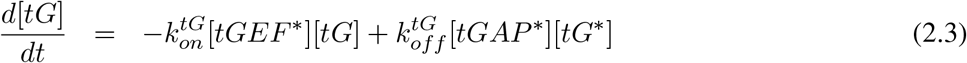

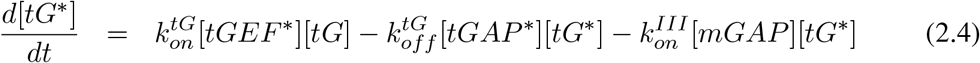

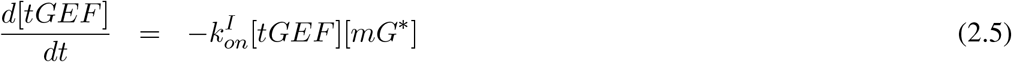

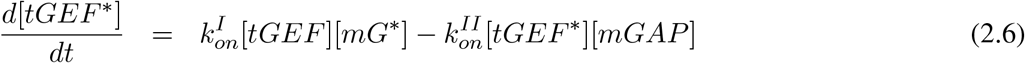

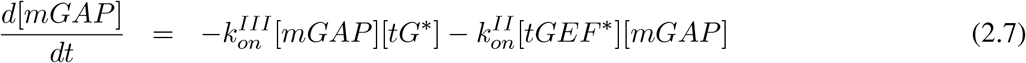

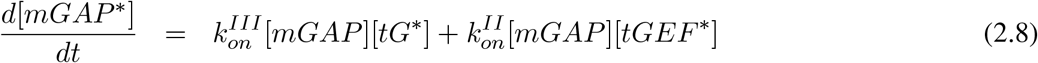

where the *k*’s represent the reaction rate parameter for each reaction rate. Since all of reactions rates are second order, the *k*’s have units of 1*/*[*µM* · *s*]

To complete the system definition, all model components must have nonnegative initial conditions. We also assume that the concentrations of [*mGEF* ^∗^] and [*tGAP* ^∗^] are constant and nonzero in our model. In particular, if 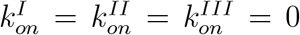, then our system describes two uncoupled GTPase switches (Fig.1B) such that each has the same dynamics of the single GTPase model proposed in [4].

### 2.3. Nondimensionalization

We introduce a nondimensional version of Eqs. 2.1 – 2.8 to reduce the number of free parameters and to obtain a new system of equations that is independent of the units of measurement. We denote 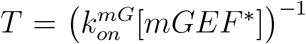 as be the characteristic time scale, ([*T*] = *s*). While there are many choices of time scales, this is the natural choice because it reflects the time scale of the coupling of the m- and t-GTPase switches. We define the characteristic concentration, *U*_*ζ*_ = [*mGEF* ^∗^] for *ζ* ∈ {*mG, mG*^∗^, *tG, tG*^∗^, *tGEF, tGEF* ^∗^, *mGAP, mGAP* ^∗^}, with units ([*U*_*ζ*_] = *µM*).

These characteristic quantities allow us to express the dimensionless kinetic rates as ratios between their dimensional forms and the rate of mGTPase activation 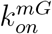. In fact, defining

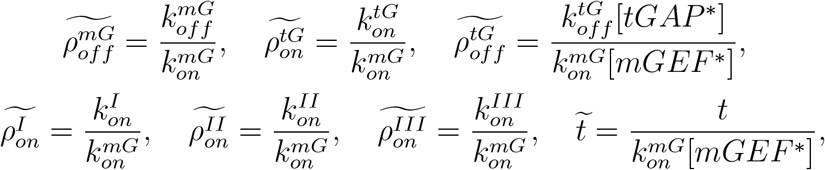

and 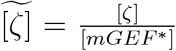 for *ζ* ∈ {*mG, mG*^∗^, *tG, tG*^∗^, *tGEF, tGEF* ^∗^, *mGAP, mGAP* ^∗^}, we drop the tildes and write the system of dimensionless equations in the following form:

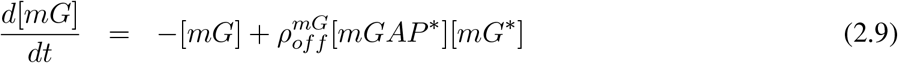

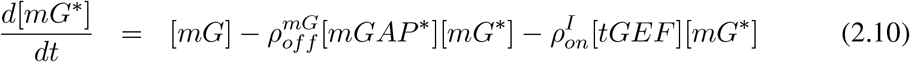

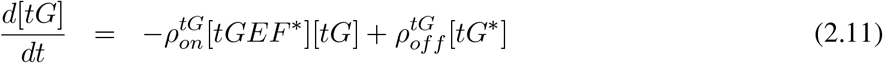

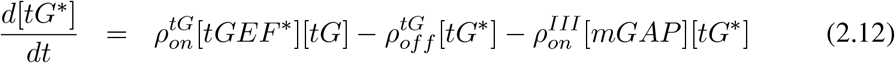

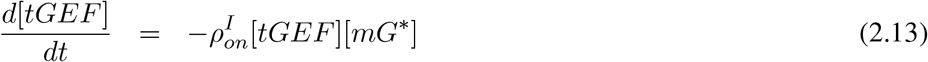

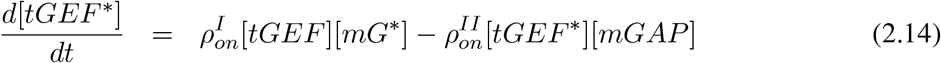

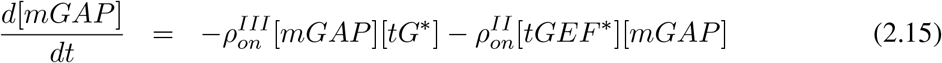

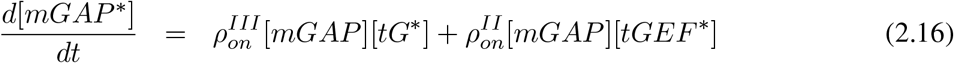

In section 3, we perform a local stability analysis of the system given by Eqs.2.9 – 2.16.

## 3. Mathematical analysis and results

In this section, we explore the role of the coupling of the two switches and feedback loops on the system dynamics. We refer to the nondimensional concentrations and rates of the system given by Eqs.2.9 – 2.16 as solely by concentrations and rates. First, it is convenient to rewrite our nondimensional ODE system in the form

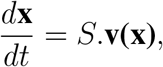

where **x** represents the vector of concentrations for the different components, *S* is the stoichiometric matrix and **v(x)** is a vector with the different reaction rates [69, 70]. Thus we define the components **x**(1) = [*mG*], **x**(2) = [*mG*^∗^], **x**(3) = [*tG*], **x**(4) = [*tG*^∗^], **x**(5) = [*tGEF*], **x**(6) = [*tGEF* ^∗^], **x**(7) = [*mGAP*], and **x**(8) = [*mGAP* ^∗^]. We also write the reaction velocities as

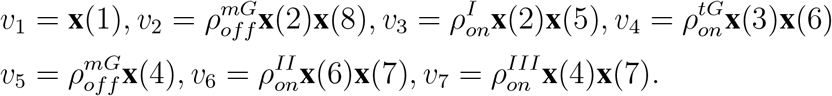

The 8 × 7 stoichiometric matrix for the system given by Eqs. 2.1 – 2.8 is then given by

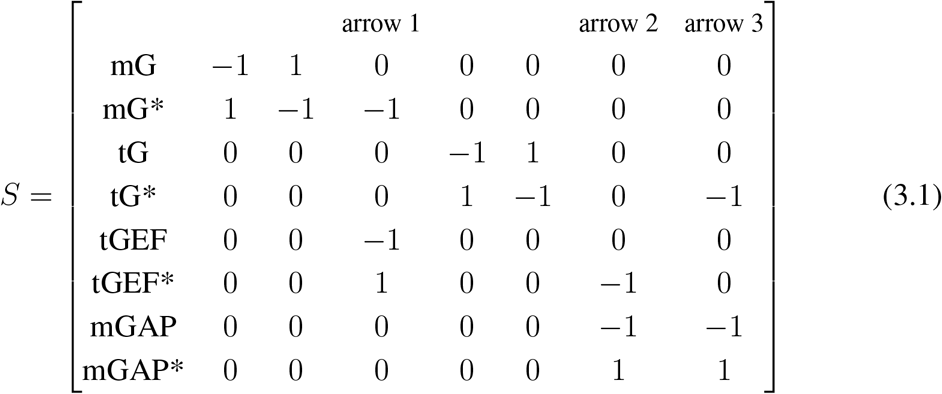

where the rows and columns of *S* (Eq. 3.1) represent the 8 components and 7 reactions, respectively. The right null space of *S* comprises the steady state flux solutions, and the left null space contains the conservation laws of the system [69]. On the other hand, the column space contains the dynamics of the time-derivatives, and the rank of S is the actual dimension of the system in which the dynamics take place. In the following subsections, we assume that 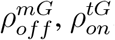, and 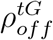 are strictly positive and we analyze Eqs. 2.9 – 2.16 when the mGTPase and tGTPase switches are coupled through: (i) A forward coupling *mG*^∗^ → *tGEF* only (arrow 1), (ii) forward coupling *mG*^∗^ → *tGEF* and feedback loop *tGEF* → *mGAP* (arrows 1 and 2) and (iii) forward coupling connection *mG*^∗^ → *tGEF* and feedback loops *tGEF* → *mGAP* and *tG*^∗^ → *mGAP* (arrows 1, 2, and 3). While mathematically, other combinations of arrows are possible, biologically, these are the only relevant combinations either for cellular function or for experimental manipulation.

### 3.1. Forward coupling connection: Recruitment of tGEF by active mGTPases (*mG*^∗^ → *tGEF*)

To analyze Eqs. 2.9 – 2.16 with the forward coupling connection only (arrow 1 in Fig.1C), we assume 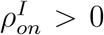 and 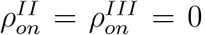, which means that the feedback loops (arrows 2 and 3) are not considered this first analysis. This represents the simple connection of the two GTPase switches, which, in cells appears to be mediated via activation–dependent coupling of mG* to tGEF [32]. In this case, the stoichiometric matrix (Eq. 3.1) is 8 × 5.

#### Conservation laws

For this particular system, the total concentrations [*tG*_*tot*_] := [*tG*] + [*tG*^∗^] and [*tGEF*_*tot*_] := [*tGEF*] + [*tGEF* ^∗^] are constant over time and are strictly positive. For this reason, it is convenient to introduce the fractions 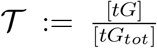 and 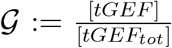 of inactive tGTPase and tGEF in the system, respectively, and let 𝒯^∗^ and G^∗^ denote the fraction of their active forms. We then use 𝒯 + 𝒯 ^∗^ = 1 and 𝒢 + 𝒢^∗^ = 1 to rewrite the system in the form

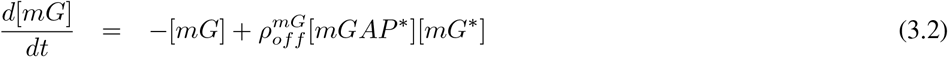

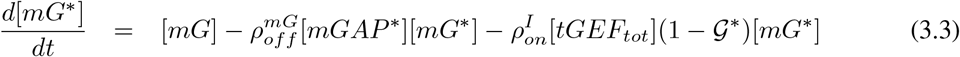

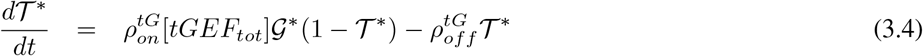

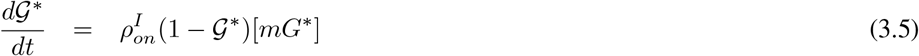

From the stoichiometric matrix (Eq. 3.1), we observe that [*mG*]+[*mG*^∗^]+[*tGEF* ^∗^] = *C*, where *C >* 0 is constant over time, and thus

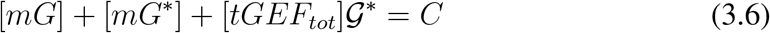

follows by the definition of 𝒢^∗^. We compute the left null space of the stoichiometric matrix and confirm the total of three conservation laws in this case. The conservation law given by Eq. 3.6 reduces the system to three unknowns, which eases the steady state and stability analysis.

#### Steady states

To find biologically plausible (nonnegative) steady states of the system given by Eqs. 3.2 – 3.6, we must find 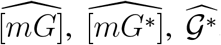, and 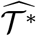 such that the time-derivatives in Eqs. 3.2–3.5 are zero and the conservation law given by Eq. 3.6 is satisfied. Therefore, we must solve the following system:

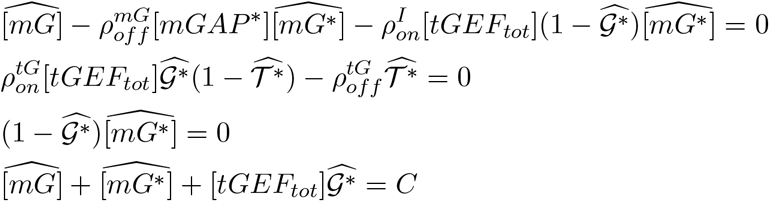

From the third equation above, we must have 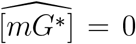 or 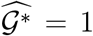. Thus we divide the steady state analysis in these two cases and summarize our results in the following proposition, whose proof can be found in the appendix A.

##### Proposition 3.1.

*The steady states* 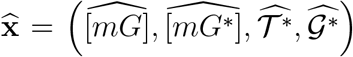 *of the system given by Eqs. 3.2 – 3.6 are given by*

- *Steady state 1:*

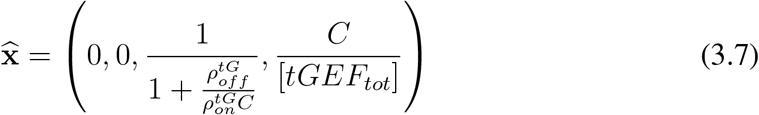 *if and only if C* ≤ [*tGEF*_*tot*_] *and*
- *Steady state 2:*

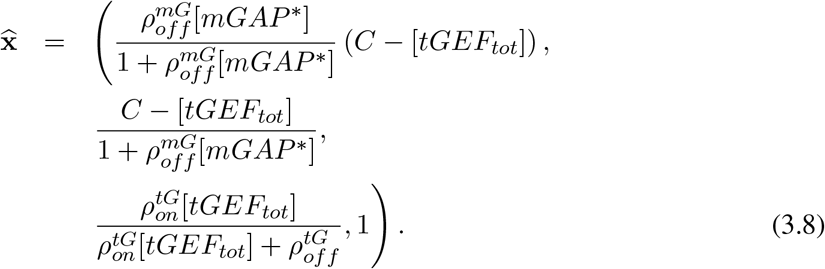

*if and only if C* ≥ [*tGEF*_*tot*_].

Given the explicit expressions for the steady states and the parameter range in which they exist, we perform a local stability analysis to determine if these states are stable or unstable under small perturbations. We adopt the classical linearization procedure based on the powerful Hartman-Grobman theorem [61, 62]. We show that the steady states are *locally asymptotically stable*, which means that any trajectory will be attracted to the steady state provided the initial condition is sufficiently close.

#### Local Stability

*Analysis*. Using that [*mG*] = *C* − [*mG*^∗^] − [*tGEF*_*tot*_]𝒢^∗^ (from Eq. 3.6) in Eqs. 3.2 – 3.5, we obtain the following three-dimensional system:

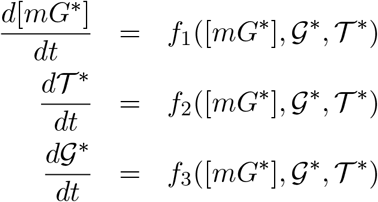

Where

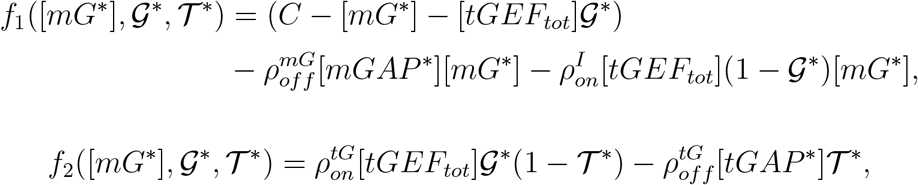

and

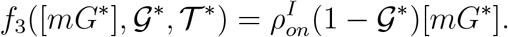

To perform the local stability analysis, we calculate the Jacobian matrix evaluated at the steady state

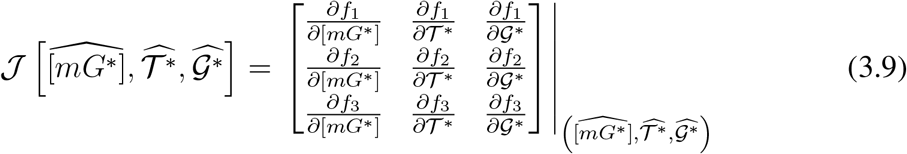

and by showing that all its eigenvalues have a negative real part, we can prove that the steady state is LAS [61], provided we further assume that the strict inequalities from Proposition 3.1 hold. This is the content of the following theorem.

##### Theorem 3.1.

*Let C be the conservation quantity from Eq. 3.6. Then*,

1. *If C <* [*tGEF*_*tot*_], *the steady state 1 (Eq. 3.7) is LAS*.
2. *If C >* [*tGEF*_*tot*_], *the steady state 2 (Eq. 3.8) is LAS*.

*Proof*. All calculations were done with MATLAB’s R2019b symbolic toolbox using the functions *jacobian* and *eig* to compute the Jacobian matrices and their eigenvalues, respectively. We proceed with the analysis of each case separately.

1. Suppose *C <* [*tGEF*_*tot*_]. As we have seen in the previous subsection, in this case the steady state is given by Eq. 3.7. The Jacobian matrix (Eq. 3.9) is given by

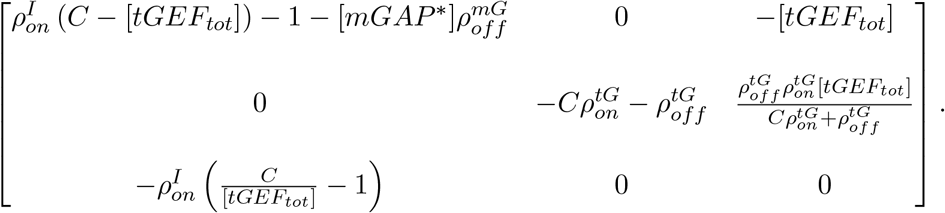

The first eigenvalue in this case is given by 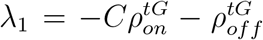 and the other two (*λ*_2_ and *λ*_3_) are such that

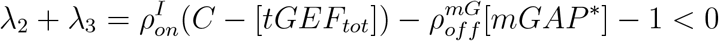

and

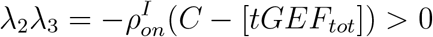

from which we conclude that *λ*_2_ and *λ*_3_ are both negative and therefore the steady state is LAS.
2. Suppose now that *C >* [*tGEF*_*tot*_]. Following our previous analysis, the steady state is given by Eq. 3.8. The Jacobian matrix in this case is given by

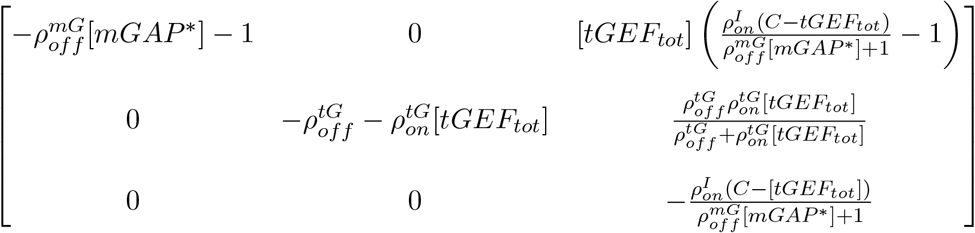

and the eigenvalues are given by 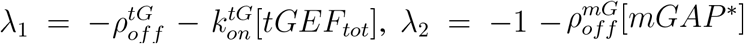 and 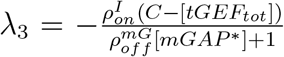, which are all negative and this completes the proof.

The inequality *C <* [*tGEF*_*tot*_] must hold for existence and local asymptotic stability to steady state 1. Recalling the definition of 𝒢^∗^ and that Eq. 3.6 holds for all times, including *t* = 0, this relationship between *C* and [*tGEF*_*tot*_] can be rewritten as [*mG*](0) + [*mG*^∗^](0) *<* [*tGEF*](0) where [*tGEF*](0) = [*tGEF*_*tot*_] − [*tGEF* ^∗^](0) is initial concentration of cytosolic tGEF that is yet to be recruited by mG* to the membranes. Similarly, the steady state 2 will exist when [*mG*](0) + [*mG*^∗^](0) *>* [*tGEF*](0). In this case, the reduced system will converge to a state where some distribution of mG, mG*, tG, tG* are present (Eq. 3.8), given sufficiently close initial and steady state concentration values. Thus, the existence of the steady states depends only on the initial concentrations and not on any kinetic parameters.

Fig 1(E) illustrates the two possible steady states (gray-colored “1” in the 2 × 2 table) promoted by the coupling connection. steady state 1 can be interpreted as a configuration where the copy numbers of both active and inactive mGTPase are low, while the tGTPase copy numbers remain high. On the other hand, in steady state 2 both m- and tGTPases have high copy numbers in both their active and inactive forms. Our results suggest that the coupling from mG to tGEF, which initiates the coupling between the two G protein switches, can drive the system to two possible configurations depending on the cellular concentrations of total mG and tGEF. If the initial tGEF is larger than the total mG, the coupling connection will result in a significant decrease of the total mG and result in the activation of a fraction of the tGEF (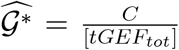 in Eq. 3.7). On the contrary, if the initial tGEF is less than the total mG, then the available tGEF will be fully engaged (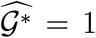 in Eq. 3.8) and there will be a residual mG concentration in the system. We conclude that the initial difference between the copy numbers of total mG and tGEF (a cytosolic protein that is recruited to the membrane by mG*) is the main factor that will determine the steady state of the coupled GTPase switches.

### 3.2 Coupled switches with feedback loop *tGEF* → *mGAP* : Recruitment of tGEF by active mGTPases and tGEF colocalization with mGAP

We analyze the case where the feedback loop *tGEF* → *mGAP* (arrow 2 in Fig.1C) is added to the coupled system with the forward connection. In cells, this feedback loop represents a tGEF* colocalization with mGAP on Golgi membranes that facilitates the recruitment of GAP proteins Fig.1A, [32]. To analyze the effects of Arrows 1 and 2 solely, we thus assume 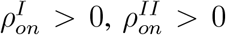 and 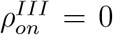. The model equations are thus given by the following system:

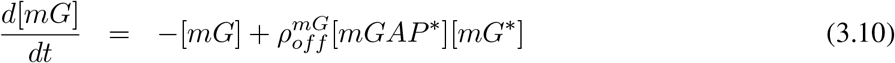

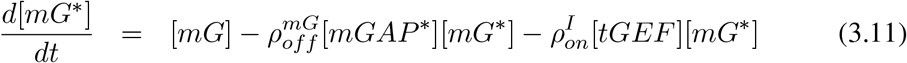

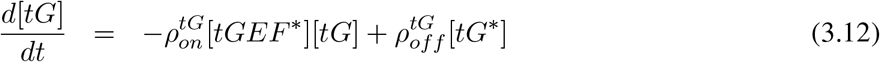

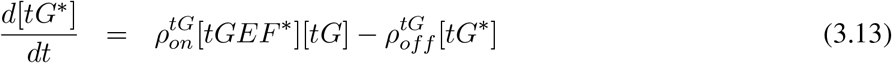

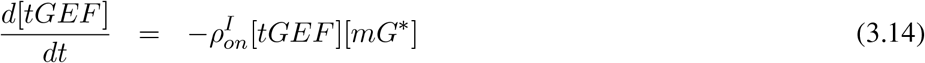

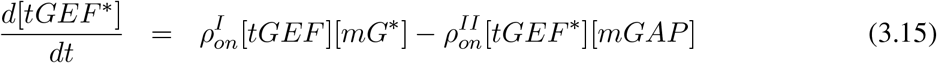

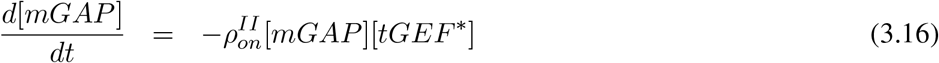

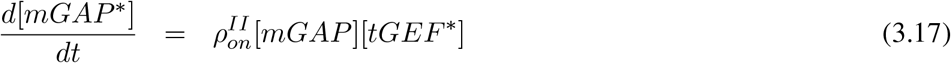

As in Section 3.1, we first analyze the conservation laws of this particular system. In this case, the stoichiometric matrix (Eq. 3.1) is 8 × 6.

#### Conservation Laws

We begin by observing that the total amount of tGTPase is conserved in this system. Thus we may use the fraction 𝒯 ^∗^ as in Section 3.1 and that is the first conservarion law. The total amount of mGAP is also conserved, as we sum Eqs. 3.16 and 3.17. We can then write

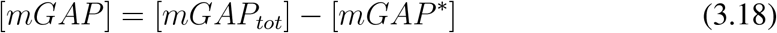

and substitute the above expression for [*mGAP*] in Eqs. 3.15 and 3.17. We choose to keep the concentrations of mGAP as a variable for notational simplicity and do not define its fraction. Summing Eqs. 3.10, 3.11, 3.15, and 3.17, and integrating over time, we get

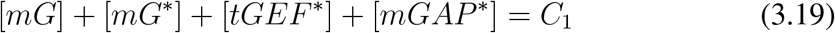

where *C*_1_ ≥ 0 is constant over time. Moreover, Eqs. 3.14, 3.15, and 3.17 when summed and integrated give

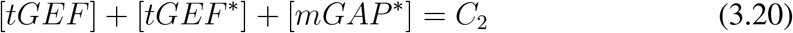

for *C*_2_ ≥ 0 also constant. We compute the left null space of the stoichiometric matrix (Eq. 3.1) and confirm a total of four conservation laws in this case. The reduced system is given by the following equations:

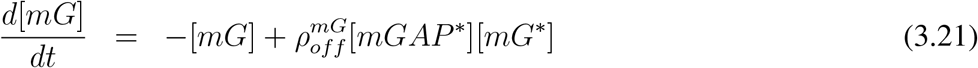

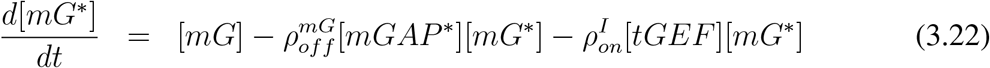

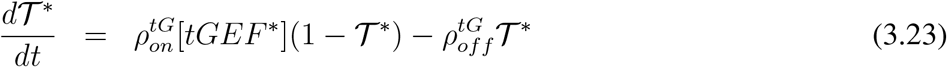

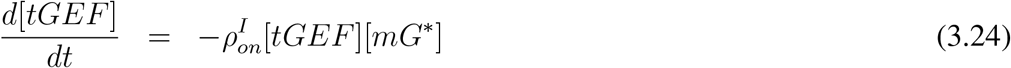

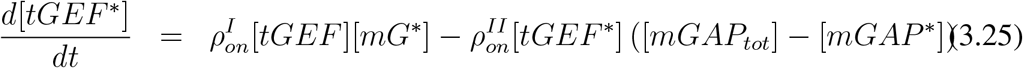

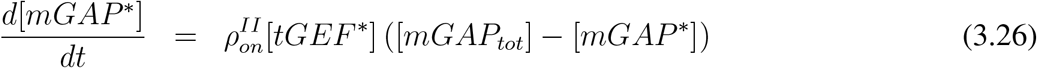

with the conservation laws given by Eqs. 3.19 and 3.20. Next, we obtain the steady states of the system.

#### Steady states and local stability analysis

To find the steady states, we must find non-negative solutions of the following system:

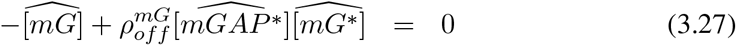

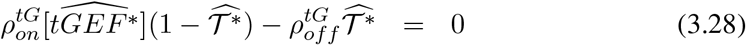

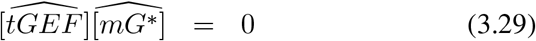

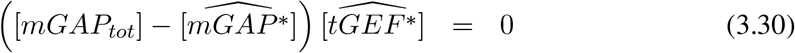

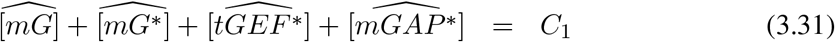

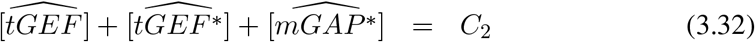

From Eq. 3.29, we must have 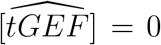 or 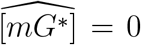. Moreover, from Eq. 3.30, 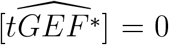 or 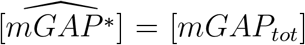 and thus we have four possible combinations to analyze.

We study each case separately and obtain the necessary and sufficient inequalities involving the parameters *C*_1_, *C*_2_, and [*mGAP*_*tot*_] that ensure the existence of each steady state. As in Section 3.1, we also show that the steady states are LAS provided the strict inequalities are satisfied. We summarize our analysis in the following theorem, whose proof can be found in Appendix B.

##### Theorem 3.2.

*The steady states*

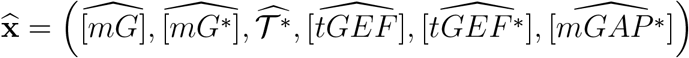

*of the system given by Eqs. 3.19 - 3.26 are given by*

- *Steady state 1:*

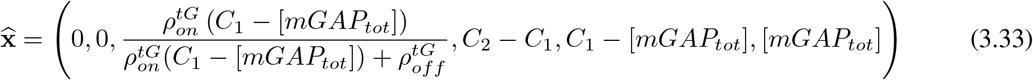

*if and only if C*_2_ ≥ *C*_1_ *and C*_1_ ≥ [*mGAP*_*tot*_]. *The steady state is LAS if C*_2_ *> C*_1_ *and C*_1_ *>* [*mGAP*_*tot*_].
- *Steady state 2:*

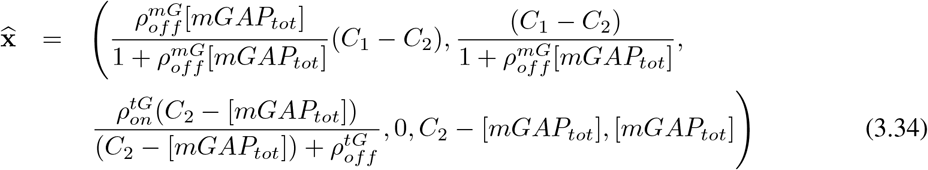

*if and only if C*_1_ ≥ *C*_2_ *and C*_2_ ≥ [*mGAP*_*tot*_]. *The steady state is LAS if C*_1_ *> C*_2_ *and C*_2_ *>* [*mGAP*_*tot*_].
- *Steady state 3:*

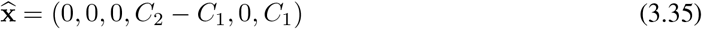

*if and only if C*_2_ ≥ *C*_1_ *and C*_1_ ≤ [*mGAP*_*tot*_]. *The steady state is LAS if C*_2_ *> C*_1_ *and C*_1_ *<* [*mGAP*_*tot*_].
- *Steady state 4:*

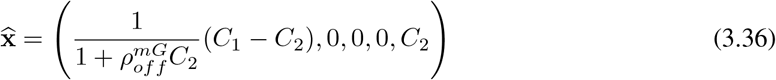

*if and only if C*_1_ ≥ *C*_2_ *and C*_2_ ≤ [*mGAP*_*tot*_]. *The steady state is LAS if C*_1_ *> C*_2_ *and C*_2_ *<* [*mGAP*_*tot*_].

Recalling the definitions of *C*_1_ and *C*_2_ and the fact that Eqs. 3.19 and 3.20 hold at all times, including at *t* = 0, we can write *C*_1_ = [*mG*](0) + [*mG*^∗^](0) + [*tGEF* ^∗^](0) + [*mGAP* ^∗^](0) and *C*_2_ = [*tGEF*](0) + [*tGEF* ^∗^](0) + [*mGAP* ^∗^](0). In this way, from the inequalities obtained in Theorem 3.2 for *C*_1_ and *C*_2_, we obtain relationships among the initial conditions of the original system (Eqs. 3.10 – 3.17) that are associated with each one of the four steady states.

For the existence and local asymptotic stability of steady state 1 (Eq. 3.33), where mG and mG* have zero concentration values, the inequalities *C*_2_ *> C*_1_ and *C*_1_ *>* [*mGAP*_*tot*_] must hold. The first inequality can be written as [*mG*](0) + [*mG*^∗^](0) *<* [*tGEF*](0), which was obtained in Section 3.1 as the existence condition for the steady state with no mG and mG* (Eq. 3.7). On the other hand, the inequality *C*_1_ *>* [*mGAP*_*tot*_] can be written as [*mG*](0) + [*mG*^∗^](0) + [*tGEF* ^∗^](0) *>* [*mGAP*](0), where [*mGAP*](0) is the initial concentration of cytosolic mGAP that is yet to be recruited by tGEF* to the membranes. Therefore, two conditions guarantee the existence of steady state 1: (1) The total amount of mG protein must be initially less than the concentration of tGEF and (2) The sum of the concentrations of total mG protein and tGEF* must be initially higher than the concentration of mGAP. If both conditions hold, then Theorem 3.2 ensures that steady state 1 will emerge and the reduced system (Eqs. 3.21 – 3.26 along with Eqs. 3.19 and 3.20) will converge to the steady state 1, provided the initial and steady state concentration values are sufficiently close.

A similar analysis holds for steady states 2, 3 and 4. For simplicity, we present the required initial conditions for each steady state without repeating the conclusions that follows from Theorem 3.2. For steady state 2 (Eq. 3.34), where mG, mG*, tG, and tG* are present, the inequalities *C*_1_ *> C*_2_ and *C*_2_ *>* [*mGAP*_*tot*_] become [*mG*](0) + [*mG*^∗^](0) *>* [*tGEF*](0) and [*tGEF*](0) + [*tGEF* ^∗^](0) *>* [*mGAP*](0), respectively. Hence, the total amount of mG protein must be initially higher than the concentration of tGEF and the total amount of tGEF must be initially higher than concentration of mGAP. For steady state 3 (Eq. 3.35), where mG and mG* have zero concentration values and the tGTPase is fully inactivated, the inequalities *C*_2_ *> C*_1_ and *C*_1_ *<* [*mGAP*_*tot*_] become [*mG*](0) + [*mG*^∗^](0) *<* [*tGEF*](0) and [*mG*](0) + [*mG*^∗^](0) + [*tGEF* ^∗^](0) *<* [*mGAP*](0), respectively. Hence, the total amount of mG protein must be initially less than the concentration of tGEF and the sum of the concentrations of total mG protein and tGEF* must be initially less than the concentration of mGAP. For steady state 4 (Eq. 3.36), where mG and mG* are present and tG* concentration is zero, the inequalities *C*_1_ *> C*_2_ and *C*_2_ *<* [*mGAP*_*tot*_] become [*mG*](0) + [*mG*^∗^](0) *>* [*tGEF*](0) and [*tGEF*](0) + [*tGEF* ^∗^](0) *<* [*mGAP*](0), respectively. Hence, the total amount of mG protein must be initially higher than the concentration of tGEF and the total amount of tGEF must be initially less than the concentration of mGAP. In this case, the existence of the steady states also depends only on the initial concentrations and not on any kinetic parameters.

Fig 1(E) illustrates the four possible steady states (gray-colored “1+2” in the 2 × 2 table) promoted by the coupled switches in the presence of the feedback loop *tGEF* → *mGAP*. Steady states 1 and 2 have the same interpretation of the two steady states obtained in Section 3.1. On the other hand, steady states 3 and 4 were obtained through the sole contribution of the feedback loop *tGEF* → *mGAP*. These states share the common feature of having tGTPase fully inactivated. However, steady state 3 can be interpreted as a configuration where the copy numbers of mG and mG* are low, while in steady state 4, these copy numbers are high.

### 3.3. Coupled switches with feedback loops *tGEF* → *mGAP* and *tG*^∗^ → *mGAP* : Recruitment of tGEF by active mGTPases, tGEF colocalization with mGAP, and activation of mGAP by active tGTPases

We analyze the case where the feedback loop *tG*^∗^ → *mGAP* (arrow 3 in Fig.1C) is added to the coupled system in addition to the feedback loop *tGEF* → *mGAP*. This connection represents the release of free *Gβγ* promoting mGAP activation. We analyze the full system given by Eqs. 2.9 – 2.16 in the case where 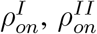, and 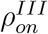 are strictly positive. In particular, we obtain the conservation laws and four 1-parameter steady state families. We also obtain the necessary conditions for the conserved quantities that guarantee the existence of each steady state family.

#### Conservation Laws

As in Section 3.2, we observe that the total amount of mGAP is constant over time, so Eq. 3.18 still holds. On the other hand, the total tGTPase follows a new conservation law that we derive here. Summing Eqs. 2.9 – 2.12, 2.14, and 2.16 and integrating over time, we have

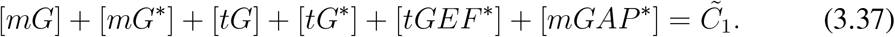

Summing Eqs. 2.11 – 2.14 and Eq. 2.16 and integrating over time, we obtain

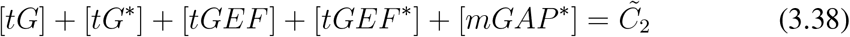

where 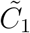 and 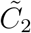 must be nonnegative constants. We compute the left null space of the stoichiometric matrix (Eq. 3.1) and confirm the total of three conservation laws, which are given by Eqs. 3.18, 3.37, and 3.38. These equations reduce Eqs. 2.9 – 2.16 to a five-dimensional system, whose steady states can be obtained.

#### Steady states

We compute the steady states of the system when the time derivatives in Eqs. 2.9 – 2.16 are equal to zero. Removing the linearly dependent equations, the problem reduces to finding the nonnegative solutions of the following system:

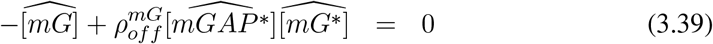

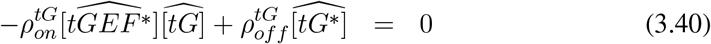

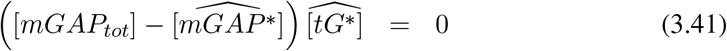

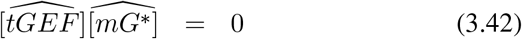

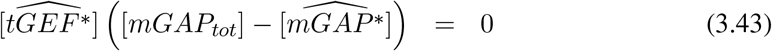

along with the conservation laws given by Eqs. 3.18, 3.37, and 3.38.

Eq. 3.40 gives 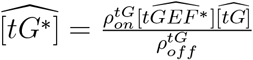 and Eq. 3.41 then becomes

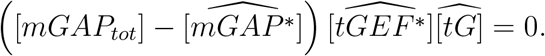

From Eq. 3.43, we conclude that 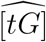 can be any nonnegative real number satisfying Eqs. 3.37 and 3.38. We define 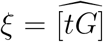 and characterize four *ξ*-dependent families of steady states similarly as we did in Section 3.2. We summarize our results in the following theorem, whose proof can be found in the appendix C.

##### Theorem 3.3.

*The ξ-dependent families of steady states*

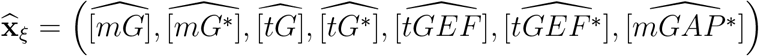

*of the system given by Eqs. 3.39 – 3.43 with the conservation laws given by Eqs. 3.18, 3.37, and 3.38 are given by*

- *Family 1:*

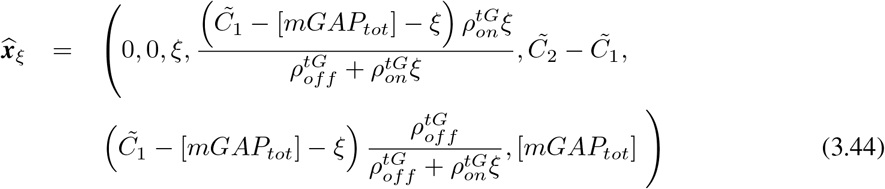

*only if* 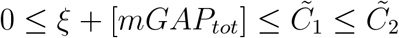.
- *Family 2:*

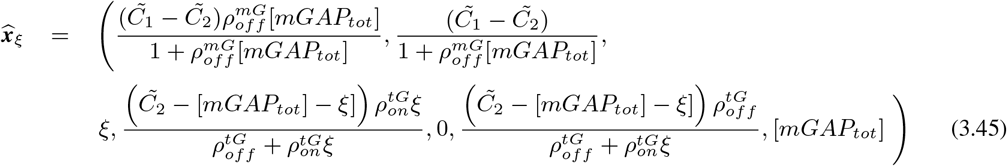

*only if* 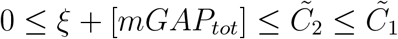.
- *Family 3:*

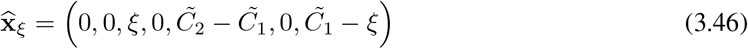

*only if* 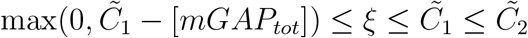.
- *Family 4:*

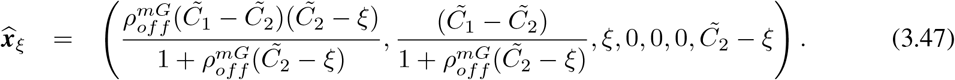

*only if* 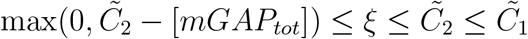.

Recalling the definitions of 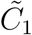 and 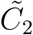 and the fact that Eqs. 3.37 and 3.38 hold at all times, including *t* = 0, we can infer necessary relationships among the initial conditions for each steady state family.The inequality 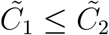 can be rewritten as [*mG*](0) + [*mG*^∗^](0) ≤ [*tGEF*](0) is necessary for the emergence of Family 1 (Eq. 3.44) whith zero mG and mG* values, which can be interpreted as a scenario in which nearly all the available mG proteins are activated to mG*, and that nearly all the mG* species have successfully engaged with the available tGEFs, thereby maximally recruiting tGEF on the membranes. For Family 1, 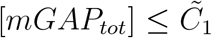 also holds and can be written as [*mGAP*](0) ≤ [*mG*](0) + [*mG*^∗^](0) + [*tG*](0) + [*tG*^∗^](0) + [*tGEF* ^∗^](0), where [*mGAP*](0) is the initial concentrations of cytosolic mGAP that is yet to be recruited by tGEF* and tG* to the membranes. Therefore, two initial conditions are necessary for the existence of Family 1: The total amount of mG protein must be initially less than the concentration of tGEF and (2) The summed concentrations of total mG, total tG and tGEF* must be initially higher than the concentration of mGAP. Finally, the inequality 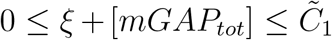 can be written as 0 ≤ *ξ* ≤ [*mG*](0)+[*mG*^∗^](0)+[*tG*](0)+[*tG*^∗^](0)+[*tGEF* ^∗^](0)−[*mGAP*](0). Remarkably, we conclude that the initial *balance* between the summed concentrations of total mG, total tG, tGEF* and the available mGAP is the upper bound for the tG concentration, which completely characterizes the necessary conditions for the emergence of Family 1.

A similar analysis can be done for Families 2, 3, and 4. For the existence of Family 2 (Eq. 3.45), where mG, mG* tG, tG* are present (when *ξ >* 0), the inequalities 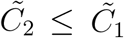 and 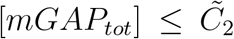 must hold and can be rewritten as [*mG*](0) + [*mG*^∗^](0) ≥ [*tGEF*](0) and [*mGAP*](0) ≤ [*tG*](0) + [*tG*^∗^](0) + [*tGEF*](0)[*tGEF* ^∗^](0). Hence, the total amount of mG protein must be initially higher than the concentration of tGEF and the summed concentrations of total tG and total tGEF proteins must be initially higher than the concentration of mGAP. Finally, the inequality 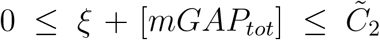 indicates that initial balance between the summed concentrations of total tG, total tGEF and the available mGAP is the upper bound for the tG concentrations. For Family 3 (Eq. 3.46, where mG and mG* have zero concentration values and the tGTPase is fully inactivated, the inequality 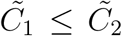 becomes [*mG*](0) + [*mG*^∗^](0) ≤ [*tGEF*](0). As for Family 1, the total amount of mG protein must be initially less than the concentration of tGEF. Moreover, from 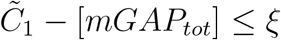, the initial balance between the summed concentrations of total mG, total tG, tGEF* and the available mGAP is the lower bound for the tG concentration. For Family 4 (Eq. 3.47), where mG and mG* are present and tG* concentration is zero, 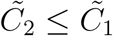 becomes [*mG*](0) + [*mG*^∗^](0) ≥ [*tGEF*](0). As for Family, 2 the total amount of mG protein must be initially higher than the concentration of tGEF. Moreover, from 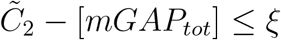, the initial balance between the summed concentrations of total tG, total tGEF and the available mGAP is the lower bound for the tG concentrations. As noted in the previous subsections, the existence of the steady states depends only on the initial concentrations and not on any kinetic parameters.

Fig 1(E) illustrates the four Families (gray-colored “1+2+3” in the 2 × 2 table) promoted by the coupling connection *mG*^∗^ → *tGEF* and the feedback loops *tGEF* → *mGAP* and *tG*^∗^ → *mGAP*. Families 1 and 2 have a similar interpretation of the steady states 1 and 2 obtained in Section 3.1 and Section 3.2. On the other hand, Families 3 and 4 were obtained through contributions of the feedback loop *tG*^∗^ → *mGAP*. These states share the common feature of having tGTPase fully inactivated. As for steady states 3 and 4, Family 3 can be interpreted as a configuration where the copy numbers of mG and mG* are low, while in Family 4, those copy numbers are high.

### 3.4. Numerical Simulations

To complete our mathematical analysis, we numerically investigate the range of initial conditions in which the trajectories of the original system (Eqs. 2.1 – 2.8) converge to the different steady states. In particular, we illustrate the so-called *basins of attraction* [71] of the steady states, considering the same combination of connections between the two GTPase switches from §3.1 – §3.3.

In Table 2, we describe each parameter of the system with the corresponding values that we used in our simulations. All ODEs were numerically solved in MATLAB R2018a with the function *ode15s*. The MATLAB codes can be downloaded in the link: https://github.com/Rangamani-Lab/BMB_Matlab_codes.git

**Table 2.** Table of parameters and initial conditions

| Parameter | Meaning | Value/range | Unit |
| --- | --- | --- | --- |
| $k_{on}^{mG}$ | activation rate of the mGTPase | 3 (Figs 2,3,4, 5 A,B,C); $\mathcal{N}(30,1)$ (Fig 5 D,E,F) | $(s.\mu M)^{-1}$ |
| $k_{off}^{mG}$ | deactivation rate of the mGTPase | 1 (Figs 2,3,4, 5 A,B,C); $\mathcal{N}(10,1)$ (Fig 5 D,E,F) | $(s.\mu M)^{-1}$ |
| $k_{on}^{tG}$ | activation rate of the tGTPase | 3 (Figs 2,3,4, 5 A,B,C); $\mathcal{N}(30,1)$ (Fig 5 D,E,F) | $(s.\mu M)^{-1}$ |
| $k_{off}^{tG}$ | deactivation rate of the tGTPase | 1 (Figs 2,3,4, 5 A,B,C); $\mathcal{N}(10,1)$ (Fig 5 D,E,F) | $(s.\mu M)^{-1}$ |
| $k_{on}^I$ | activation rate of tGEF through forward coupling | 0 or 3 (Figs 2,3,4, 5 A,B,C); 0 or $\mathcal{N}(30,1)$ (Fig 5 D,E,F) | $(s.\mu M)^{-1}$ |
| $k_{on}^{II}$ | activation rate of mGAP through feedback loop (arrow 2) | 0 or 3 (Figs 2,3,4, 5 A,B,C); 0 or $\mathcal{N}(30,1)$ (Fig 5 D,E,F) | $(s.\mu M)^{-1}$ |
| $k_{on}^{III}$ | activation rate of mGAP through feedback loop (arrow 3) | 0 or 3 (Figs 2,3,4, 5 A,B,C); 0 or $\mathcal{N}(30,1)$ (Fig 5 D,E,F) | $(s.\mu M)^{-1}$ |
| $[mGEF^*]$ | concentration of active mGEF | 1 (Figs 2,3,4,5) | $\mu M$ |
| $[tGAP^*]$ | concentration of active tGAP | 1 (Figs 2,3,4,5) | $\mu M$ |
| $[tG_{tot}]$ | Total concentration of tGTPase | 10 (Figs 2,3,4); 5 (Fig 5) | $\mu M$ |
| $[tGEF_{tot}]$ | Total concentration of tGEF | 10 (Figs 2,3,4); 5 (Fig 5) | $\mu M$ |
| $[mG](0)$ | initial concentration of inactive mGTPase | [0,10] (Figs 2,3,4); 0 (Fig 5) | $\mu M$ |
| $[mG^*](0)$ | initial concentration of active mGTPase | 0 (Figs 2,3,4); [0,10] (Fig 5) | $\mu M$ |
| $[tG](0)$ | initial concentration of inactive tGTPase | 5 (Figs 2,3,4,5) | $\mu M$ |
| $[tG^*](0)$ | initial concentration of active tGTPase | 5 (Figs 2,3,4); 0 (Fig 5) | $\mu M$ |
| $[tGEF](0)$ | initial concentration of inactive mGTPase | 5 (Figs 2,3,4,5) | $\mu M$ |
| $[tGEF^*](0)$ | initial concentration of active tGEF | 5 (Figs 2,3,4) or 0 (Fig 5) | $\mu M$ |
| $[mGAP](0)$ | initial concentration of inactive mGAP | [0,20] (Figs 3,4); [0,12] (Fig 5) | $\mu M$ |
| $[mGAP^*](0)$ | initial concentration of inactive mGAP | 1 (Figs 2,3,4); [0,12] (Fig 5) | $\mu M$ |

In Fig.2, we explore the case where the two GTPase switches are coupled through the coupling connection *mG*^∗^ → *tGEF* (Fig.2A). We color the trajectories of the system according to the comparison between the initial conditions [*mG*](0) + [*mG*^∗^](0) and [*tGEF* ^∗^](0) from the steady state analysis in Section 3.1. For fixed [*mG*^∗^](0) and [*tGEF*](0) values, we consider [*mG*](0) ranging from 0 to 10 *µM* and therefore [*mG*](0) + [*mG*^∗^](0) can be less of higher than [*tGEF*](0) (blue or red-colored lines and dots). For all simulations, we plot the trajectories of the system until equilibrium is reached. If [*mG*](0) + [*mG*^∗^](0) *<* [*tGEF*](0), the system converges to a state where no active mGTPase exists (blue colored trajectories in Figs.2B and 2C). On the other hand, if [*mG*](0)+[*mG*^∗^](0) *>* [*tGEF*](0), the system converges a state where the concentration of the active and inactive mGTPase are positive at the final time (red-colored trajectories). To visualize these results in terms of dose-response curves, in Fig.2D we plot the final-state values of [*mG*_*tot*_] and 𝒢^∗^ (denoted by s.s) as a function of [*mG*](0). The trajectories in the 𝒯 ^∗^ × [*mG*_*tot*_] plane are shown in Fig.2E. We observe a detail showing that 𝒯 ^∗^ reaches a fixed final value around 0.97 when [*mG*](0) + [*mG*^∗^](0) *>* [*tGEF*](0) (see magnified view). We observe that the trajectories converge to steady states that agree with the local stability results from §Section 3.1. This suggests that the conditions [*mG*](0) + [*mG*^∗^](0) *<* [*tGEF*](0) and [*mG*](0) + [*mG*^∗^](0) *>* [*tGEF*](0) are not only valid in a neighborhood of the steady states, but also hold for other initial values satisfying those inequalities.

**Figure 2.**
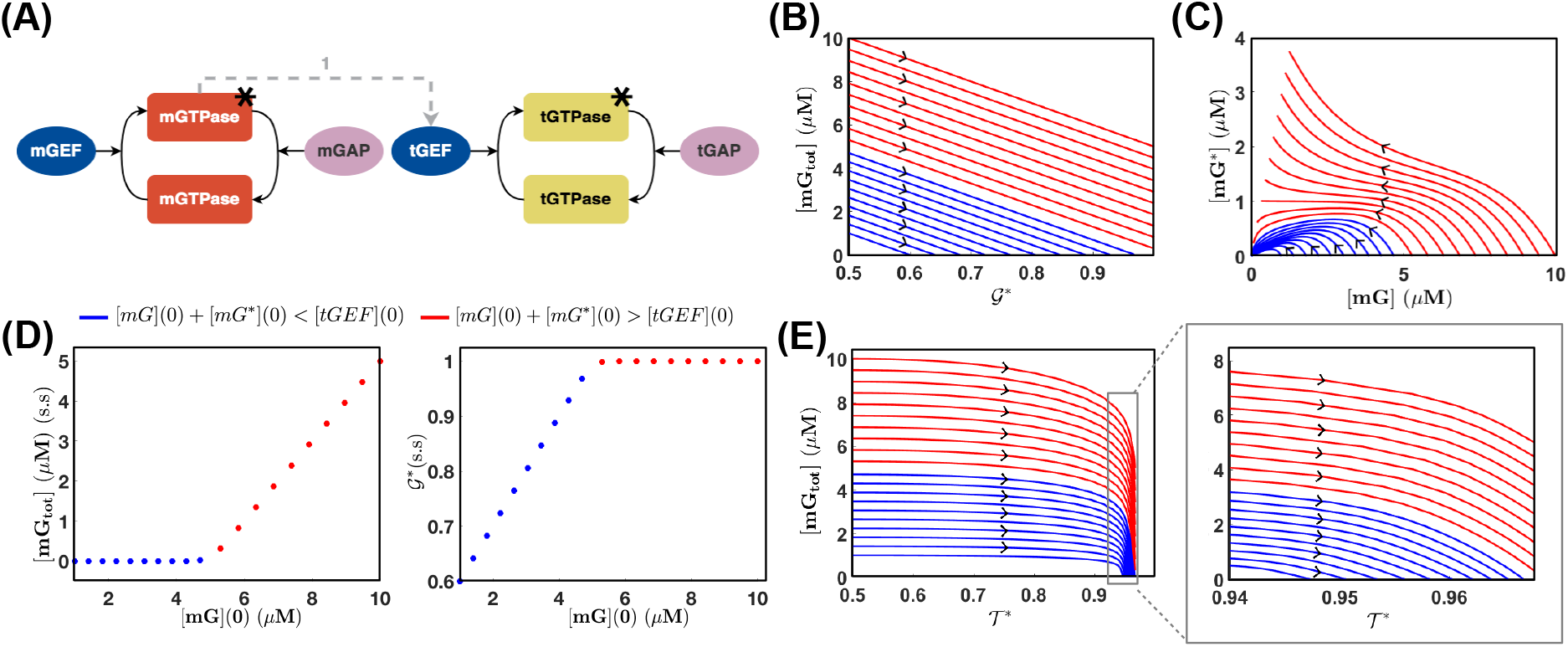
Trajectories of the system and steady states (arrow 1) (A) Schematics with the coupled GTPase switches and a coupling connection *mG*^∗^ → *tGEF*, represented by arrow 1. (B) [*mG*](0) was changed from 0 to 10 *µM* and the trajectories of the system were calculated until equilibrium was reached. In the 𝒢^∗^ × [*mG*_*tot*_] plane, a linear relationship emerges. The black arrows indicate the direction of time. If [*mG*](0) *>* 5*µM*, the system converges to a final-state where the concentrations of the active and inactive mGTPase are nonzero. On the other hand, when [*mG*](0) *<* 5*µM*, the trajectories converge a final-state with no mGTPase exists (blue colored lines). (C) Trajectories of the active ([*mG*]) vs inactive mGTPase ([mG]) for [*mG*](0). (D) Dose response curves show the steady states (denoted by s.s) for the total mGTPase concentration and fraction of active tGEF (G) depending on [*mG*](0) in the two different scenarios. (E) The dynamics in the 𝒯 × [*mG*_*tot*_] plane. Parameter values: [*mG*^∗^](0) = 0*µM*, 𝒯 ^∗^(0) = 0.5, 𝒢^∗^(0) = 0.5, [*mGAP* ^∗^] = 1*µM*, [*mGEF* ^∗^] = 1*µM, k*_*on*_ = 3(*s.µM*)^−1^, *k*_*off*_ = 1(*s.µM*)^−1^, [*tGAP* ^∗^] = 1*µM*, [*tGEF*_*tot*_] = 10*µM*, [*tG*_*tot*_] = 10*µM*. Simulation time: 5*s* for panels B and E, 50*s* for panels C, D, F, and G. Numerical simulations were performed using the solver ode15s in Matlab R2018a. All parameters were arbitrarily chosen only to illustrate the dynamic features of the model.

Fig.3 illustrates the dynamics of the system when the feedback loop *tGEF* → *mGAP* (Arrow 2) is added to the coupling connection (Fig.3A). In Fig.3B, we plot several [*tGEF* ^∗^] trajectories starting at [*tGEF* ^∗^] = 5*µM* for different [*mG*](0) and [*mGAP*](0) values. The resulting rich variety of curves indicate the sensitivity of the system to these initial conditions. In Fig.3C, different dose-response curves are generated to show the steady state tGEF* values. If [*mGAP*](0) = 0*µM* (blue and red dots), only the coupling connection affects the system, since mGAP cannot be activated by tGEF*. When [*mGAP*](0) = 1 (green squares), a similar steady state profile emerges, with [*tGEF* ^∗^] s.s increasing for [*mG*](0) ≤ 5*µM* and remaining constant [*mG*](0) *>* 5. When [*mGAP*](0) = 8*µM*, [*tGEF* ^∗^] is zero for [*mG*](0) *<* 2*µM* and increases until [*mG*](0) *<* 5*µM*. For [*mG*](0) *>* 5, the steady state achieves its maximum value slightly above [*tGEF* ^∗^] *>* 2. Finally when [*mGAP*](0) = 11*µM*, *tGEF* ^∗^ becomes fully recruited by mGAP and the [*tGEF* ^∗^] s.s is zero for all [*mG*](0) values. In Fig.3D, we scan the space of initial amounts of mG and mGAP. When [*mGAP*](0) *>* 10*µM*, the [*tGEF* ^∗^] s.s is zero, while for [*mGAP*](0) *<* 10*µM* is becomes nonzero and dependent of [*mG*](0). In Figs. 3 (E), (F), and (G), we analyze the tG* concentration values and obtain similar results.

**Figure 3.**
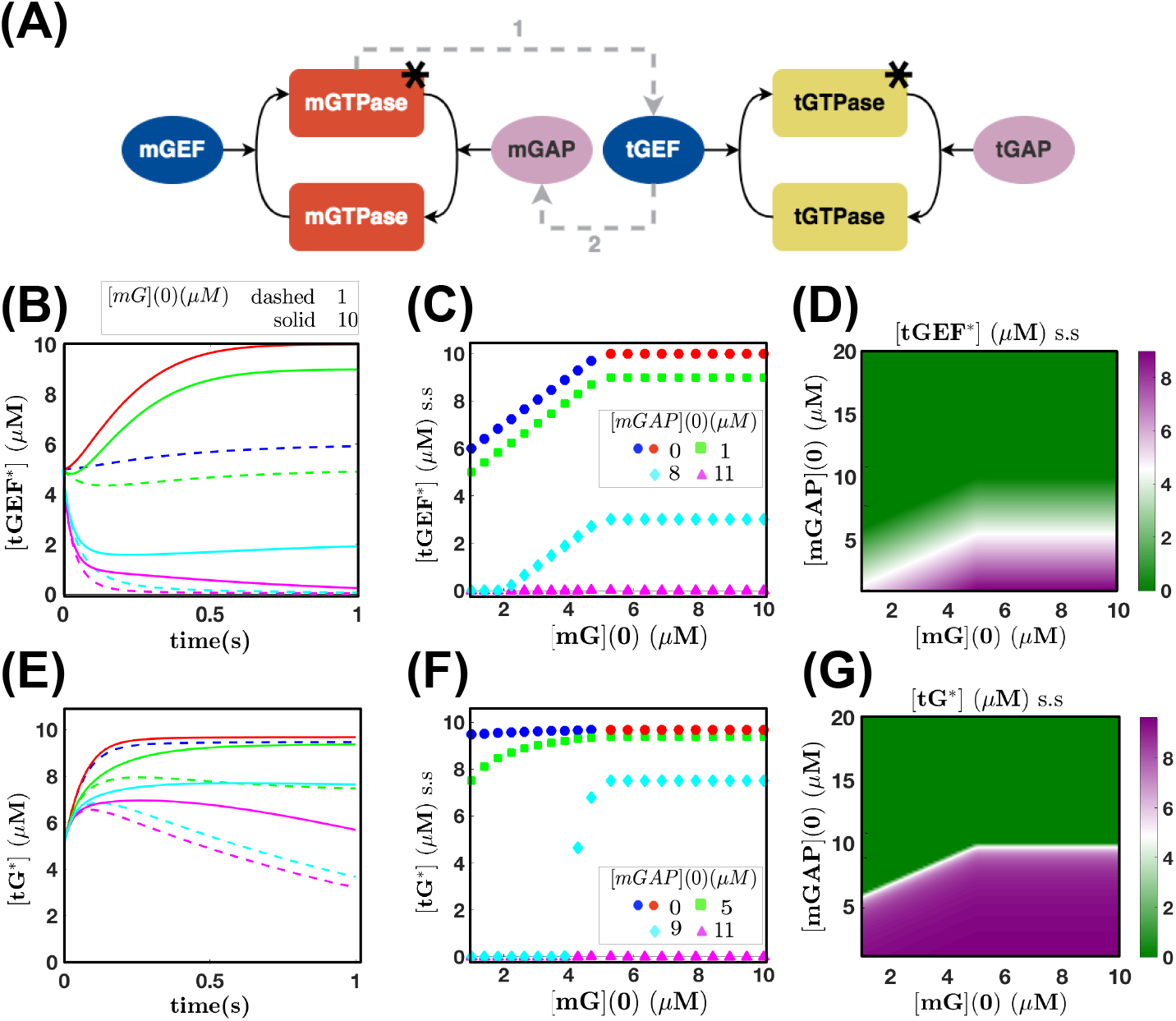
Trajectories of the system and steady states (s.s) (arrows 1 and 2). (A) Schematics of the coupled GTPases with the coupling connection *mG*^∗^ → *tGEF* (arrow 1) and the feedback loop *tGEF* → *mGAP* (arrow 2). (B) [*tGEF* ^∗^] trajectories for [*mGAP*](0) = 0, 1, 8, and 11 *µM*. For each [*mGAP*](0) value, we plot two curves for [*mG*](0) = 1 (dashed) and 10 *µM* (solid) (C) Dose response curves show [*tGEF* ^∗^] s.s when [*mG*](0) ranges from 0 to 10 *µM*. If [*mGAP*](0) = 0*µM* (blue and red dots), there will be no mGAP activation and therefore no effects of the feedback. For [*mGAP*](0) *>* 0*µM*, the feedback becomes effective and generate different [*tGEF* ^∗^] responses. (D) Colormap for [*tGEF* ^∗^] s.s concentrations for a range of [*mG*](0) and [*mGAP*](0) values. A sharp decrease on [*tGEF* ^∗^] occurs when [*mGAP*](0) ≥ 10*µM*. When [*mGAP*](0) *<* 10*µM*, the [*tGEF* ^∗^] s.s depend on [*mG*](0). (E) [*tG*^∗^] trajectories for [*mGAP*](0) = 0, 5, 9, and 11 *µM* and same [*mG*](0). (F) Dose response curves for [*tG*^∗^] s.s depend on [*mGAP*](0). (G) Colormap for [*tG*^∗^] s.s.; lower tG* concentrations result from higher [*mGAP*](0) values, since tGEF* is recruited for mGAP activation. Parameter values: *k*_*on*_ = 3(*s.µMs*)^−1^, *k*_*o*_*ff* = 1(*s.µM*)^−1^, [*mG*^∗^](0) = 0*µM*, [*tGEF*_*tot*_](0) = 10*µM*, [*tGEF* ^∗^](0) = 5*µM*, 𝒯 ∗(0) = 0.5, [*tG*_*tot*_] = 10*µM*, [*mGAP* ^∗^](0) = 1*µM*, [*tGAP* ^∗^](0) = 1*µM*, [*mGEF* ^∗^] = 1*µM*. Simulation times: 5*s* (B and E) and 50*s* (C, D, F, and G). Numerical simulations were performed using the solver ode23s in Matlab R2018a. All parameters were arbitrarily chosen only to illustrate the dynamic features of the model.

Fig.4 illustrates the dynamics of the system when the feedback loops *tGEF* → *mGAP* and *tG*^∗^ → *mGAP* are added to the coupling connection (Fig.4A). In Fig.3B, we plot several [*tGEF* ^∗^] trajectories starting at [*tGEF* ^∗^] = 5*µM* for different [*mG*](0) and [*mGAP*](0) values. In Fig.3C, different dose-response curves are generated to show the steady state tGEF* values. As in the previous case with only one feedback loop, if [*mGAP*](0) = 0*µM* (blue and red dots), mGAP cannot be activated by tGEF*. When [*mGAP*](0) = 1 (green squares), a similar steady state profile emerges, with [*tGEF* ^∗^] s.s increasing for [*mG*](0) ≤ 5*µM* and remaining constant [*mG*](0) *>* 5. When [*mGAP*](0) = 8 and 11 *µM*, [*tGEF* ^∗^] increases until [*mG*](0) *<* 5*µM*. For [*mG*](0) *>* 5, the steady state achieves its maximum value. In Fig.4D, we scan the space of initial amounts of mG and mGAP and we observe a more graded response in comparison with Fig.3. In Figs. 4 (E), (F), and (G), we analyze the tG* concentration values and obtain similar results.

**Figure 4.**
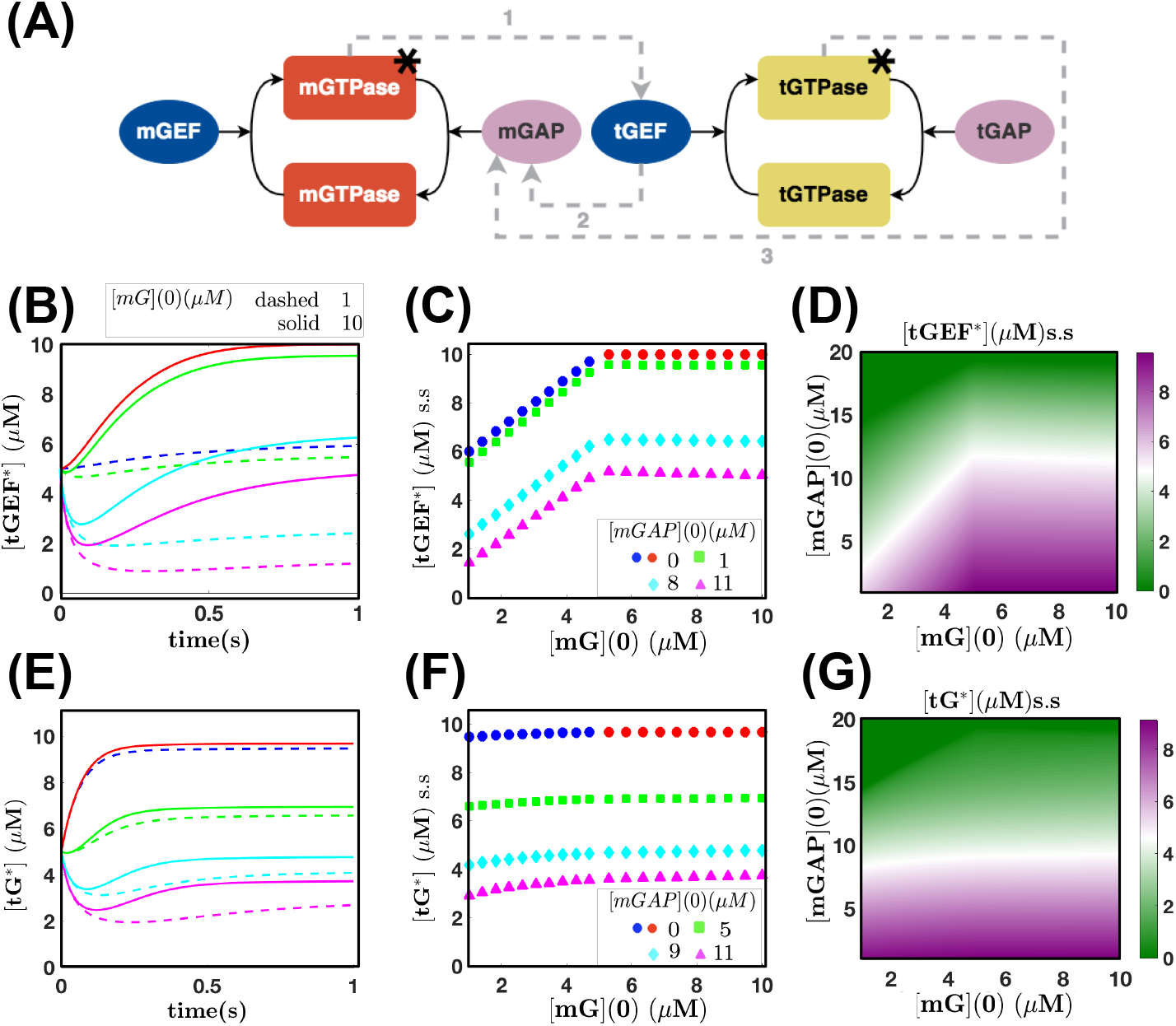
Trajectories of the system and steady states (s.s) (arrows 1, 2, and 3). (A) Schematics of the coupled GTPases with the coupling connection *mG*^∗^ → *tGEF* (arrow 1) and the feedback loops *tGEF* → *mGAP* (arrow 2) and *tG*^∗^ → *mGAP* (arrow 3). (B) [*tGEF* ^∗^] trajectories for [*mGAP*](0) = 0, 1, 8, and 11 *µM*. For each [*mGAP*](0) value, we plot two curves for [*mG*](0) = 1 and 10 *µM*. (C) Dose response curves show [*tGEF* ^∗^] s.s when [*mG*](0) ranges from 0 to 10 *µM*. If [*mGAP*](0) = 0*µM* (blue and red dots), there will be no mGAP activation and therefore no effects of the feedback loops. For [*mGAP*](0) *>* 0*µM*, the feedback becomes effective and generate different [*tGEF* ^∗^] responses. (D) Colormap for [*tGEF* ^∗^] s.s concentrations for a range of [*mG*](0) and [*mGAP*](0) values. A more graded decrease on [*tGEF* ^∗^] occurs when [*mGAP*](0) ≥ 10*µM* in comparison with Fig.3D. (E) [*tG*^∗^] trajectories for [*mGAP*](0) = 0, 5, 9, and 11 *µM* and same [*mG*](0). (F) Dose response curves for [*tG*^∗^] s.s depend on [*mGAP*](0) and does not change significantly as [*mG*](0) increases. (G) Colormap for [*tG*^∗^] s.s.; lower tG* concentrations result from higher [*mGAP*](0) values, since tGEF* and tG* are recruited for mGAP activation. Parameter values: *k*_*on*_ = 3(*s.µMs*)^−1^, *k*_*o*_*ff* = 1(*s.µM*)^−1^, [*mG*^∗^](0) = 0*µM*, [*tGEF*_*tot*_](0) = 10*µM*, [*tGEF* ^∗^](0) = 5*µM*, T ∗(0) = 0.5, [*tG*_*tot*_] = 10*µM*, [*mGAP* ^∗^](0) = 1*µM*, [*tGAP* ^∗^](0) = 1*µM*, [*mGEF* ^∗^] = 1*µM*. Simulation times: 5*s* (B and E) and 50*s* (C, D, F, and G). Numerical simulations were performed using the solver ode23s in Matlab R2018a. All parameters were arbitrarily chosen only to illustrate the dynamic features of the model.

In Fig.5, we investigate the space of initial conditions for mG* and mGAP* in which the system converges to the different steady states. Fig.5A shows the simplest system where the two GTPase switches are connected by the coupling *mG*^∗^ → *tGEF*. Two steady states are obtained depending on the initial amount of mG*. For [*mG*^∗^](0) *<* [*tGEF*](0) − [*mGAP* ^∗^](0) = 5*µM*, the trajectories converge to steady state 1 with no mG and mG* concentrations. On the other hand, for [*mG*^∗^](0) *>* [*tGEF*](0) − [*mGAP* ^∗^](0) = 5*µM*, then the system achieves the steady state 2 with non zero concentrations of both m and t-GTPase. Fig.5B shows the results for the coupling connection, and feedback loops *tGEF* → *mGAP* (arrows 1+2). In this particular example, the four steady states can be achieved for [*mGAP* ^∗^](0) and [*mG*^∗^](0) ranging from 0 to 12 *µM* and 0 and 10 *µM*, respectively. In the vertical direction, the initial amount of mG* governs the transitions from steady states 3 to 4 (lower [*mGAP* ^∗^](0)) and 1 to 2 (higher [*mGAP* ^∗^](0)). In both steady states 2 and 4 (Eqs. 3.34 and 3.36), the concentrations of mGTPase are nonzero. Therefore, we predict that an increase of initial concentration of mG* would favor the emergence of these two steady states. In the horizontal direction, when the initial amount of mGAP* increases, the available mGAP (inactive) decreases as we set the total mGAP as 12 *µM*, which reduces the effects of the feedback loops and thus facilitates the emergence of the steady states 1 and 2 where the concentrations of tGTPase are nonzero.

Fig.5C shows a similar colormap for the system with both feedback loops *tGEF* → *mGAP* and *tG*^∗^ → *mGAP*. It is worth noticing the expansion of the basin of attraction of Families 1 and 2 compared to Fig.5A, while the basin of Families 3 and 4 shrinks. Remarkably, in both Figs. 5B and 5C, there is a critical point (represented by a black cross) of intersection of the four basins of attraction. In this case, disturbances in the initial conditions around that intersection point can drive the system to different steady states. Thus, while coupling the two GTPase switches with a forward arrow only gives two possible steady states, the negative feedback afforded by arrows 2 and 3 give rise to a larger range of possibilities. Additionally, the existence of a critical point emerges in the presence of the negative feedback suggesting a rich phase space for this coupled system. Finally, in Figs 5(D-F),we sample the *k*_*on*_’s from a normal distribution with mean 30(*s.µM*)^−1^ and standard deviation 1(*s.µM*)^−1^ and new *k*_*off*_ ‘s from a normal distribution with mean 10(*s.µM*)^−1^ and standard deviation 1(*s.µM*)^−1^. We note that the system behavior does not change for changes in kinetic parameters. By doing so, we illustrate how the basins of attraction remain the same, given distinct reaction rates with a different order of magnitude.

**Figure 5.**
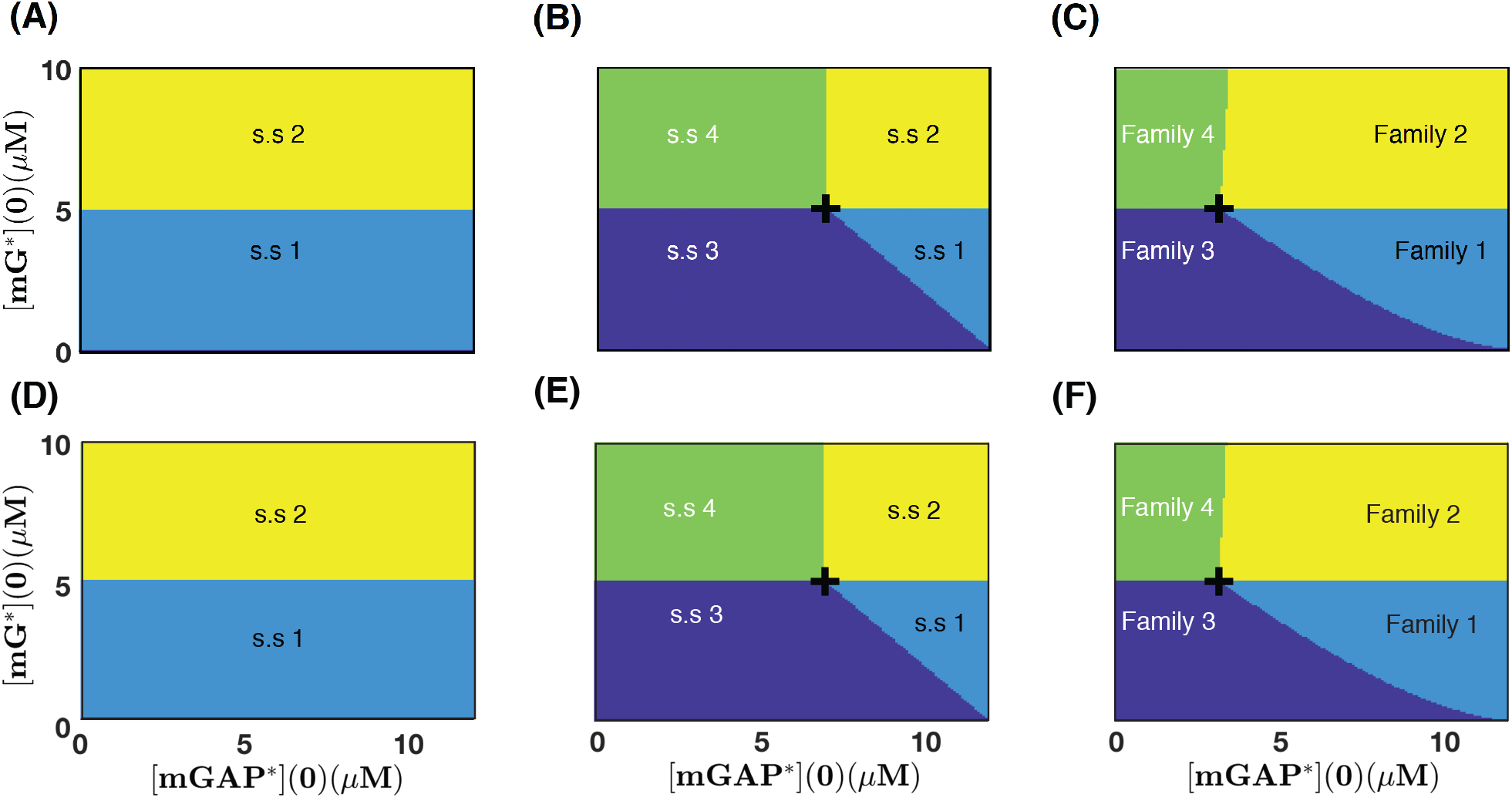
Basins of Attraction – dependency on [*mG*^∗^(0)] and [*mGAP* ^∗^(0)] (A) The two steady states of the system with coupling connection (Section 3.1) are only driven by changes in the initial amount of mG* (B) When the coupling connection and both feedback loops *tGEF* → *mGAP* are considered, we observe the emergence of four regions (green,yellow, dark blue and light blue colored) corresponding to the four steady states from Section 3.2 (C) A similar result was found when we analyzed the system with the coupling connection and both feedback loops *tGEF* → *mGAP* and *tG*^∗^ → *mGAP*. A black cross indicates a critical point at the intersection of the four basins of attraction. (D), (E), and (F) The basins of attraction remain unaltered when we consider distinct activation/deactivation rates of different orders of magnitude. Parameter values for panels (A), (B), and (C): 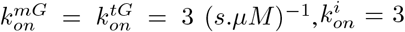 or 0 (*s.µM*)^−1^ for *i* = *I, II*, or *III*, 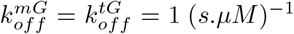. Reaction rates for panel (D) in 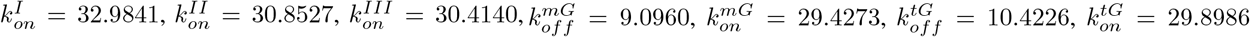, Reaction rates for panel (E) in 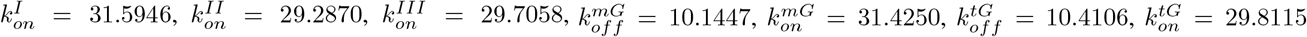 Reaction rates for panel (F) in 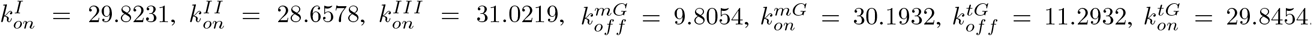. Initial conditions: [*mG*](0) = 0*µM*, [*tG*](0) = 5*µM*, [*tG*^∗^](0) = 0*µM*, [*tGEF*](0) = 5*µM*, [*tGEF* ^∗^](0) = 0*µM*, [*mGAP*](0) = 12*µM* − [*mGAP* ^∗^](0), [*tGAP* ^∗^](0) = 1*µM*, [*mGEF* ^∗^](0) = 1*µM*

## 4. Discussion

GTP-binding proteins (GTPases) regulate crucial aspects of numerous cellular events. Their ability to act as biochemical switches is essential to promote information processing within signaling networks. The two types of GTPases monomeric (m) and trimeric - have traditionally believed to function independently until recent experimental work revealed that m- and tGTPases coregulate each other in the Golgi through a functionally coupled circuit [32]. Using a simplified model of ODEs, our analyses have shown that the coupled switch gives rise to steady state configurations that cannot be achieved in systems of isolated GTPase switches. To the best of our knowledge, this is the first modeling effort that has described the stability properties of these coupled GTPase switches.

A major result from our analysis is a systematic characterization of the steady state concentrations of both m- and tGTPases, as well as their GEFs and GAPs. We show the obtained steady states in all three arrow combinations that were informed by experiments (Table 3). Remarkably, the different steady states show a variety of configurations in which both m- and tGTPase can be interpreted as having low or high concentration values. We also note that the stability properties of these steady states are independent of the choice of kinetic parameters in this model. We next interpret these different steady states in their biological context.

**Table 3.** Main results and conclusions from steady state analysis. We performed a steady state analysis of a GTPase coupled circuit that has been observed experimentally. For three biologically relevant combinations among the coupling connection and two feedback loops, we present the steady states and their interpretation. Moreover, we established the required initial conditions for the existence of the steady states. Each connection adds to the richness of the functioning of these coupled GTPase switches.

| Arrows | Name | Interpretation | Required Initial conditions | Conclusion |
| --- | --- | --- | --- | --- |
| 1 | Steady-state 1<br>Steady-state 2 | Low mG, mG*. High tG*<br>High mG, mG*. High tG* | $mG + mG^* \leq tGEF$<br>$mG + mG^* \geq tGEF$ | Feed-forward allows mG and mG* to achieve low s.s concentrations. High s.s concentrations of mG, mG*, tG, and tG* emerge if initial $mG+mG^*$ is initially higher than tGEF. |
| 1+2 | Steady-state 1<br>Steady-state 2<br>Steady-state 3<br>Steady-state 4 | Low mG, mG*. High tG,tG*<br>High mG, mG*, tG, tG*<br>Low mG, mG*, tG*<br>Low tG*. High mG,mG* | $mG + mG^* \leq tGEF$ & $mGAP \leq mG + mG^* + tGEF^*$<br>$mG + mG^* \geq tGEF$ & $mGAP \leq tGEF + tGEF^*$<br>$mG + mG^* \leq tGEF$ & $mGAP \geq mG + mG^* + tGEF^*$<br>$mG + mG^* \geq tGEF$ & $mGAP \geq tGEF + tGEF^*$ | Feed-forward and feedback $tGEF \rightarrow mGAP$ allow mG, mG*, and tG* to achieve low s.s concentrations simultaneously. High s.s concentrations of mG, mG*, tG, and tG* emerge if initials $mG+mG^*$ and $tGEF+tGEF^*$ are higher than tGEF and mGAP, resp. |
| 1+2+3 | Family 1<br>Family 2<br>Family 3<br>Family 4 | Low mG, mG*. High tG,tG*<br>High mG, mG*, tG*<br>Low mG, mG*, tG*<br>Low tG*, High mG, mG* | $mG + mG^* \leq tGEF$ & $mGAP \leq mG + mG^* + tG + tG^* + tGEF^*$<br>$mG + mG^* \geq tGEF$ & $mGAP \leq tG + tG^* + tGEF + tGEF^*$<br>$mG + mG^* \leq tGEF$<br>$mG + mG^* \geq tGEF$ | Feed-forward, feedback from tGEF to mGAP and feedback from tG* to mGAP yields four families of steady-states where the inactive tG can range within intervals. |

First and foremost, the coupling of the two switches allows for the emergence of two stable steady states. The first steady state (Eq. 3.7) has zero mG and mG* values and finite tG and tG* values. This concentration distribution of the species can be interpreted as a scenario in which nearly all the available mG proteins are activated to mG*, and that nearly all the mG* species have successfully engaged with the available tGEFs, thereby maximally recruiting tGEF on the Golgi membranes. This steady state emerges when the total amount of mG protein is less than the concentration of tGEF in cells. Similarly, the steady state 2 will emerge when the total concentration of mG is greater than tG. In this case, the reduced system will converge to steady state 2 where some finite, nonzero distribution of mG, mG*, tG, tG* are present (Eq. 3.8), given sufficiently close initial and steady state concentration values.

When we couple the connection of the two GTPase switches with the feedback loop *tGEF* → *mGAP* (arrows 1 and 2 in Fig.1 C, respectively), we obtain four steady states. Steady states 1 and 2 (Eqs. 3.33 and 3.34) are similar to the two steady states obtained in Section 3.1, although with different concentration values. On the other hand, steady states 3 and 4 (Eqs. 3.35 and Eqs. 3.36) newly emerge in the system, in which tG* attains zero concentration. This zero concentration can be interpreted as a scenario in which nearly all the available tGTPase has cycled through the GTP-cycle and is inactivated. The inequalities obtained in Theorem 3.2 for *C*_1_ and *C*_2_, allow us to obtain relationships among the initial conditions of the original system (Eqs. 3.10 – 3.17) that are associated with each one of the four steady states. Two conditions guarantee the existence of steady state 1: (1) The total amount of mG protein must be initially less than the concentration of tGEF and (2) the sum of the concentrations of total mG protein and tGEF* must be initially higher than the concentration of mGAP. Similar analysis reveals that for steady state 2 (Eq. 3.34), the total amount of mG protein must be initially higher than the concentration of tGEF and the total amount of tGEF must be initially higher than concentration of mGAP. For steady state 3 (Eq. 3.35), the total amount of mG protein must be initially less than the concentration of tGEF and the sum of the concentrations of total mG protein and tGEF* must be initially less than the concentration of mGAP. For steady state 4 (Eq. 3.36), the total amount of mG protein must be initially higher than the concentration of tGEF and the total amount of tGEF must be initially less than the concentration of mGAP.

Finally, when the coupled switches have both feedback effects on *mGAP* through Arrows 2 and 3, we obtain four families of steady states. Interestingly, the Families 1 – 4 resemble the steady states 1 – 4 from Section 3.2. Family 1 has no mG and mG* at steady state, and both tG and tG* have nonzero steady state values (similar to steady state 1). Moreover, Family 2 has both m- and tGTPases with nonzero steady states (similar to steady state 2). For Family 3, mG and mG* steady state values are zero, and the tGTPase is fully inactivated (similarly to steady state 3). Finally, Family 4 has tGTPase is fully inactivated, and both mG and mG* have nonzero steady states (similarly to steady state 4). Recalling the definitions of 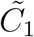 and 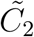 and the fact that Eqs. 3.37 and 3.38 hold at all times, including *t* = 0, we can infer necessary relationships among the initial conditions for each steady state family.

Thus, our model shows that when the m- and t-GTPase switches are coupled with a simple forward coupling (Arrow 1), there are two steady states. The addition of feedback from the t-GTPase switch to the m-GTPase switch (Arrow 2 alone or Arrows 2 and 3), expands this space to either 4 steady states or 4 families of steady states. We confirmed that all steady states obtained with a coupling connection and feedback loop *tGEF* → *mGAP* (arrow 1 or arrows 1+2 in Fig.1C) are locally asymptotically stable. However, when the two feedback loops are considered along with the coupling connection (arrows 1+2+3 in Fig.1C), the local stability analysis cannot be performed because the steady states are not isolated. Instead, we obtain four one-parameter families that depend on the amount of inactive tGTPase. At this point, further investigation would be needed to determine the behavior of the system near those steady state families. Even as we aim to develop complex models that are refined with iterative experimental validations, we note that our analysis gives insight to different steady states that emerge due to different couplings that may not exist in physiology. Such insights may become meaningful in the context of disease pathogenesis where copy numbers of each player in the network motif may change relative to each other, and do so dynamically (e.g., when responding to stress/stimuli), or disease-driving mutations alter their functions (e.g., activating and inactivating mutations in GTPases, GAPs, or GEFs).

Limitations of this study include a simplified mathematical structure of the model. Despite this simplification, we find a rich phase space for the coupled GTPase switches by analyzing the combination of network connections that have more biological meaning. Future studies could also explore the role of external stimulus, the temporal and spatial organization of these switches. While the current model is likely incomplete, it serves as a stepping stone for future adaptations that can be coupled with experimental measurements [41], including dose-response curves, response times, and noise fluctuations, as done recently in [72].

## 5. Acknowledgments

This work was supported by Air Force Office of Scientific Research (AFOSR) Multidisciplinary University Research Initiative (MURI) grant FA9550-18-1-0051 (to P. Rangamani) and the National Institute of Health (CA100768, CA238042 and AI141630 to P. Ghosh). Lucas M. Stolerman acknowledges support from the National Institute of Health (CA209891). The authors would like to acknowledge Prof. Ali Behzadan (Sacramento State University) for his careful reading and insightful comments for this work.

## Appendix A. Proof of Proposition 3.1

We must find nonnegative 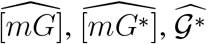, and 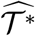 satisfying the following system:

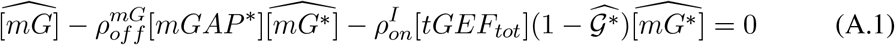

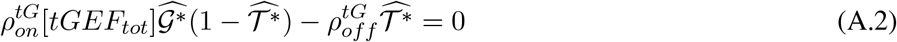

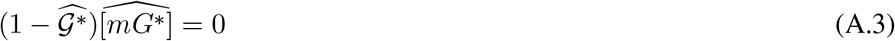

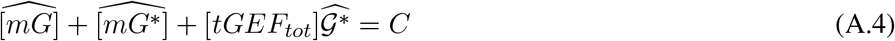

From Eq. A.3, we must have 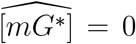 or 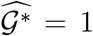. Thus we divide the steady state analysis in two cases.

Case 1: 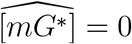.

From Eq. A.1 we must have 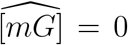 and from Eq. A.4, we obtain 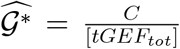. Since 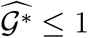 by definition, we conclude that

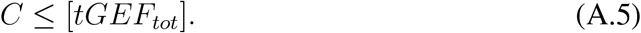

Eq. A.5 is also sufficient for 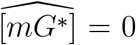. Otherwise, if *C* ≤ [*tGEF*_*tot*_] and 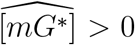, then 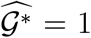 (Eq. A.3) and from Eq. A.4, we would conclude that 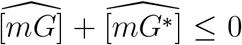, which is impossible.

Finally, by substituting 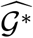 in Eq. A.2, we obtain 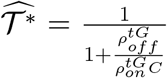 and therefore the steady state is given by

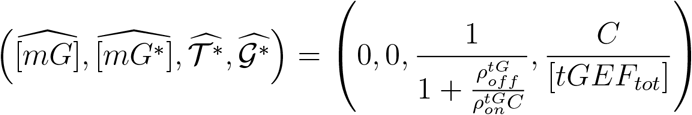

Case 2: 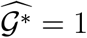

In this case, 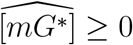 and from Eqs. A.1 and A.4,we obtain

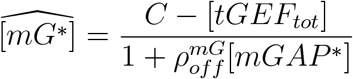

and

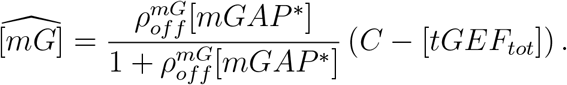

In this case, since the steady state has to be nonnegative, we must have

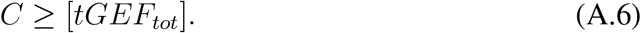

which is also sufficient for 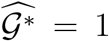. Otherwise if *C* ≥ [*tGEF*_*tot*_] and 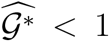, then 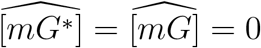 (Eqs. A.1 and A.3) and, from Eq. A.4, we would have

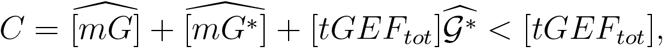

which is impossible.

Finally, by substituting 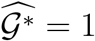 in Eq. A.2, we obtain

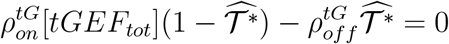

which gives 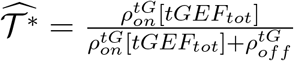 and therefore

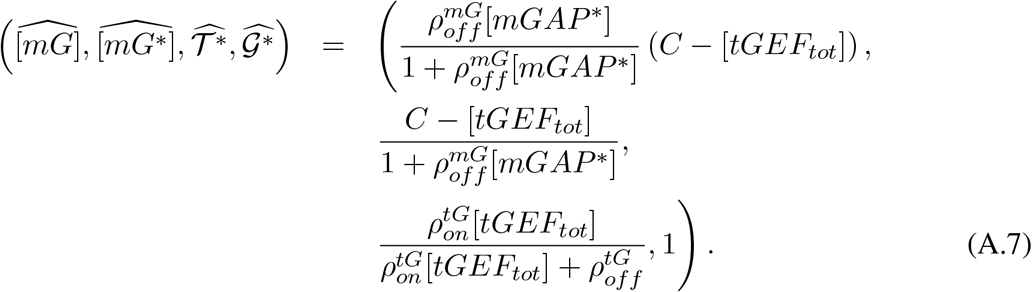

Appendix B. Proof of Theorem 3.2

We begin our proof by computing the steady states of the system, which are solutions of the algebraic system given by Eqs. 3.27–3.32. We also establish necessary and sufficient conditions involving the parameters *C*_1_, *C*_2_, and [*mGAP*_*tot*_] for the existence of each steady state. We then compute the Jacobian matrix of the system and determine the local stability of the steady state based on the classical linearization procedure [61].

### Steady states

We divide our analysis into four different cases that emerge from the preliminary inspection of the system given by Eqs. 3.27–3.32.

Case 1: 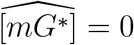 and 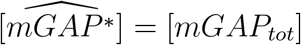.

From Eq. 3.27, we have 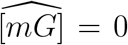 and from Eq. 3.31, 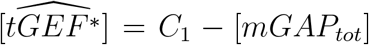. Thus *C*_1_ ≥ [*mGAP*_*tot*_] since the steady state must be nonnegative. Now Eq. 3.32 gives 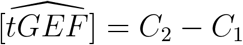 and that implies *C*_2_ ≥ *C*_1_.

Finally, Eq. 3.28 yields

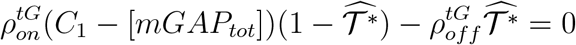

and hence

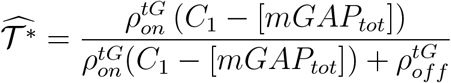

The steady state is therefore given by

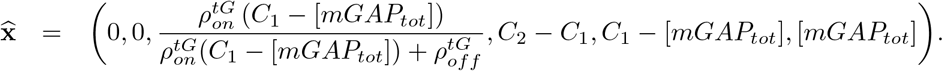

We now observe that the two parameter relations

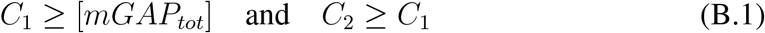

are sufficient for 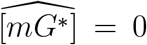 and 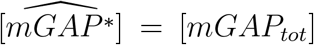. First, we observe that if *C*_2_ ≥ *C*_1_ then 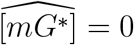. In fact, by subtracting 3.31 from Eq. 3.32, we obtain

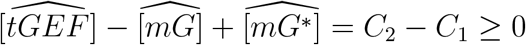

and hence 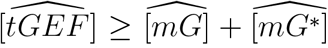. On the other hand, from Eq. 3.29, we must have 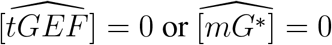. Thus if 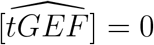 then 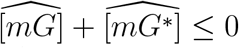 and hence the nonnegativeness of the steady state implies 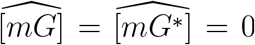. Now, Eq. 3.31 gives 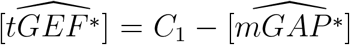 and from Eq. 3.30, we must have 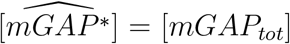 or 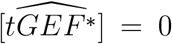. If 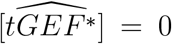, then 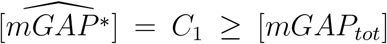 and hence 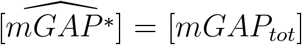. Therefore, we have shown that Eq. B.1 imply 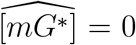 and 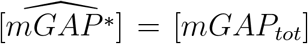. Consequently, the steady state in this case must be given by Eq. 3.33.

Case 2: 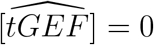 and 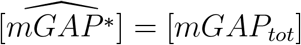

From Eq. 3.32, 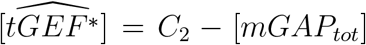 and hence [*mGAP*_*tot*_] ≤ *C*_2_. From Eq. 3.31, we must have 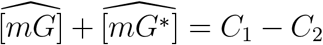 and that implies *C*_1_ ≥ *C*_2_. Now, Eq. 3.27 gives

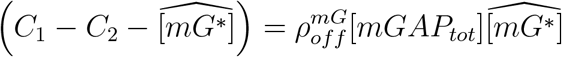

and therefore

From Eq. 3.28, we must have

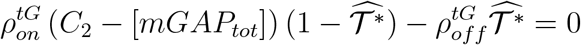

from which we obtain

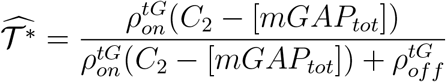

and therefore the steady state is given by

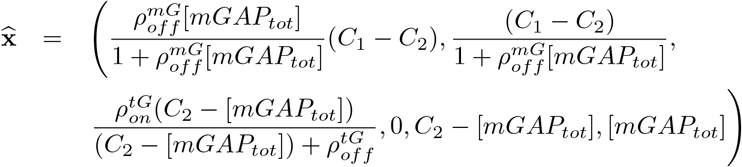

We now observe that the two parameter relations

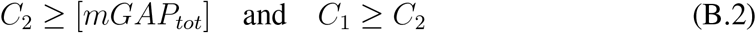

are sufficient for 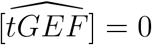 and 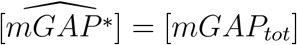.

In fact, if *C*_1_ ≥ *C*_2_ then 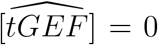 from the same argument as in Case 1. Now, Eq. 3.32 gives 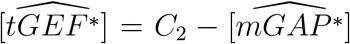] and from Eq. 3.30, we must have 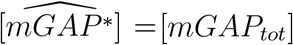 or 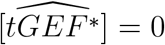. If 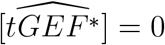 then 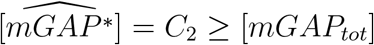 (from Eq. B.2) and thus 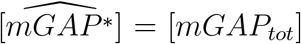. Therefore, we have shown that Eq. B.2 imply 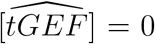 and 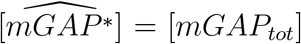. Consequently, the steady state in this case must be given by Eq. 3.34.

Case 3: 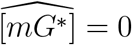 and 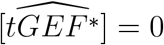.

From Eq. 3.27, we have 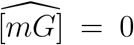 and from Eq. 3.28, we also get 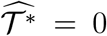 since 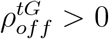. Now, Eq. 3.31 gives 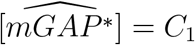 and thus we must have *C*_1_ ≤ [*mGAP*_*tot*_]. Moreover, Eq. 3.32 results in 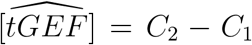 and since all steady states must be nonnegative, we obtain *C*_2_ ≥ *C*_1_. In this case, the steady state is given by

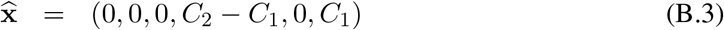

We now observe that the two parameter relations

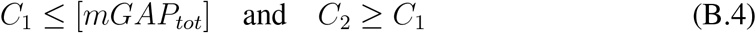

are sufficient for 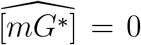 and 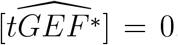. In fact, *C*_2_ ≥ *C*_1_ implies 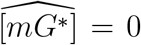 from the same argument as in Case 1.

Now, Eq. 3.31 gives 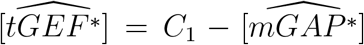 and from Eq. 3.30, we must have 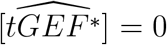 or 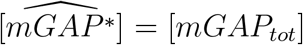. If 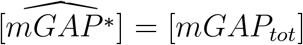, then 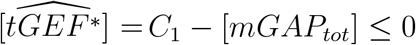 (from Eq. B.4) and thus 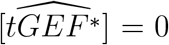. Therefore, we have shown that Eq. B.4 imply 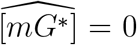 and 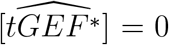. Consequently, the steady state in this case must be given by Eq. 3.35.

Case 4: 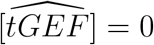 and 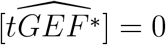

From Eq. 3.32, we obtain 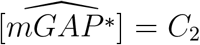 and hence *C*_2_ ≤ [*mGAP*_*tot*_]. From Eq. 3.31, we have 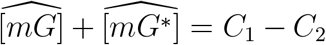 and that implies *C*_1_ ≥ *C*_2_ since the concentrations at steady state must be nonnegative. Eq. 3.27 then gives from which we obtain

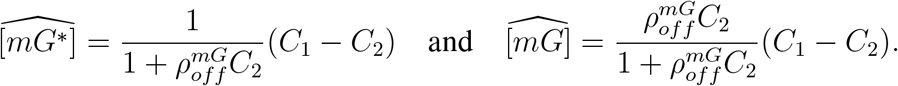

From Eq. 3.28, we have 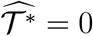 and therefore the steady state is given by

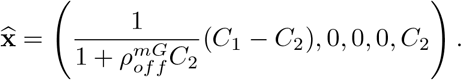

We now observe that the two parameter relations

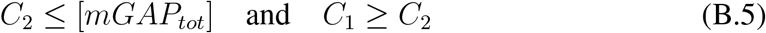

are sufficient for 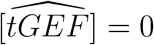 and 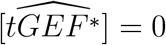. In fact, if *C*_1_ ≥ *C*_2_ then by subtracting Eq. 3.32 from Eq. 3.31, we have

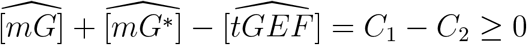

and hence 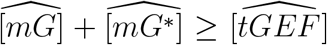. On the other hand, from Eq. 3.29, we must have 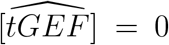 or 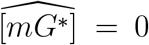. Thus if 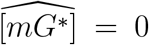 then 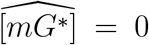 (from Eq. 3.27) and hence the nonnegativeness implies 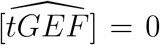. Hence we conclude that Eq. B.5 guarantee 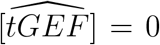.

Now, Eq. 3.32 gives 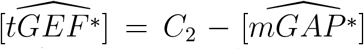 and from Eq. 3.30, we must have 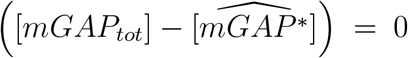 or 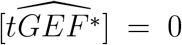. If 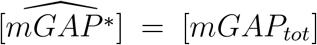 then 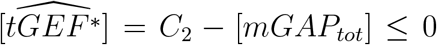 (from Eq. B.5) and thus 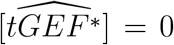. Therefore, we have shown that Eq. B.5 implies 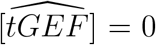 and 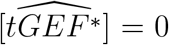. Consequently, the steady state in this case must be given by Eq. 3.36.

### Local Stability Analysis

We begin reducing the ODE system with the conservation laws given by Eqs. 3.19 and 3.20. In fact, if we write

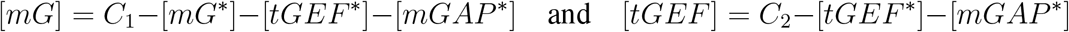

then Eqs. 3.21 – 3.26 can be written in the form

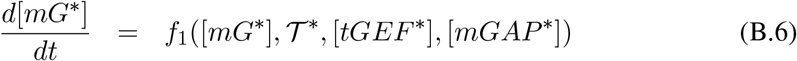

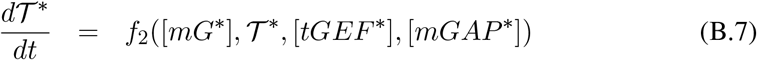

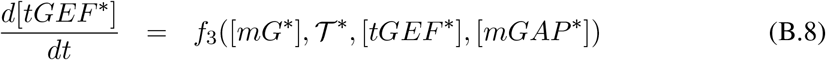

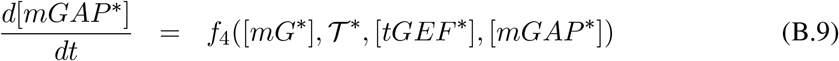

Where

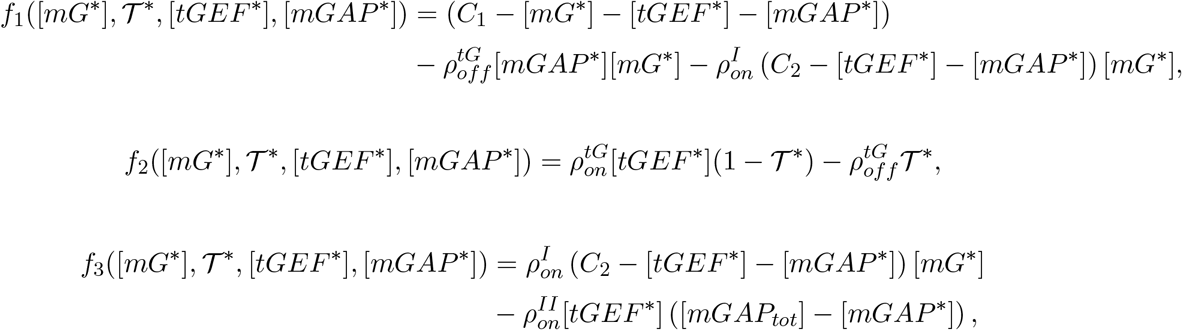

and

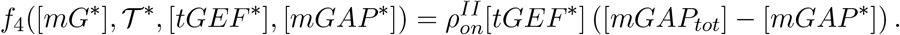

The eigenvalues of the Jacobian Matrix can be thus calculated for each one of the four steady states given by Eqs. 3.33 – 3.36. We prove that all steady states are LAS by showing that the eigenvalues of the Jacobian Matrix are all negative real numbers. We perform the calculations with MATLAB’s R2019b symbolic toolbox and analyze each case separately (see supplementary file with MATLAB codes). We analyze each case separately.

1. If *C*_1_ *>* [*mGAP*_*tot*_] and *C*_2_ *> C*_1_, the Jacobian matrix evaluated at the steady state given by Eq. 3.33 gives the eigenvalues

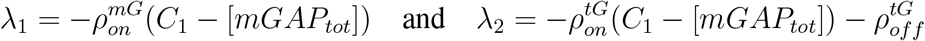

which are negative. Moreover, the other eigenvalues *λ*_3_ and *λ*_4_ are such that

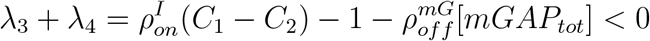

and

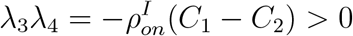

and thus *λ*_3_ and *λ*_4_ are negative and hence the steady state is LAS.
2. If *C*_2_ *>* [*mGAP*_*tot*_] and *C*_1_ *> C*_2_, the Jacobian matrix evaluated at the steady state given by Eq. 3.34 gives the eigenvalues

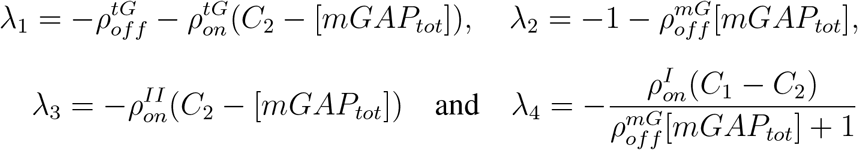

which are all negative and hence the steady state is LAS.
3. If *C*_1_ *<* [*mGAP*_*tot*_] and *C*_2_ *> C*_1_,the Jacobian matrix evaluated at the steady state given by Eq. 3.35 gives the eigenvalues

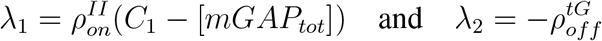

which are negative. Moreover, the other eigenvalues *λ*_3_ and *λ*_4_ are such that

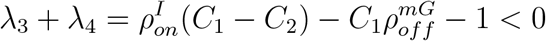

and

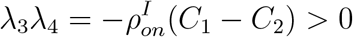

and thus *λ*_3_ and *λ*_4_ are negative and hence the steady state is LAS.
4. If *C*_2_ *<* [*mGAP*_*tot*_] and *C*_1_ *> C*_2_, the Jacobian matrix evaluated at the steady state given by Eq. 3.36 gives the eigenvalues

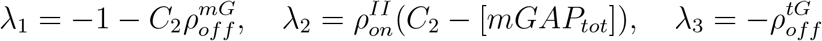

and

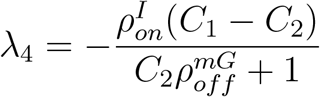

which are all negative and hence the steady state is LAS.

Appendix C. Proof of Theorem 3.3

We proceed with the steady state analysis in the same way of Theorem 3.2. We consider the same four different cases and calculate the *ξ*-dependent families of steady states, where *ξ* ≥ 0 represent the tG concentration. We also obtain necessary relationships for the conserved quantities 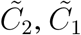, and [*mGAP*_*tot*_], as well as admissible intervals for *ξ* that guarantee the existence of nonnegative steady states.

Case 1: 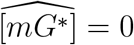 and 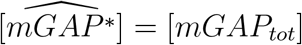.

From Eq. 3.39, we have 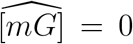 and subtracting Eq. 3.38 from Eq. 3.37, we get 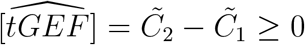 only if 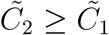. Substituting 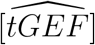 on the conservation law given by Eq. 3.38 and using Eq. 3.40 to write 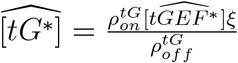, we obtain

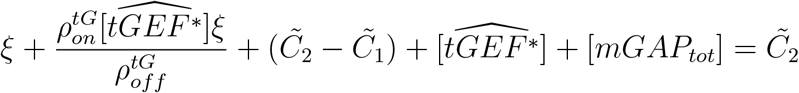

and hence

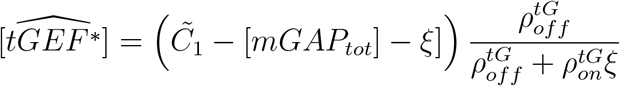

only if 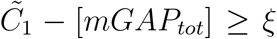. Therefore, in this case the *ξ*-dependent family of steady states is given by

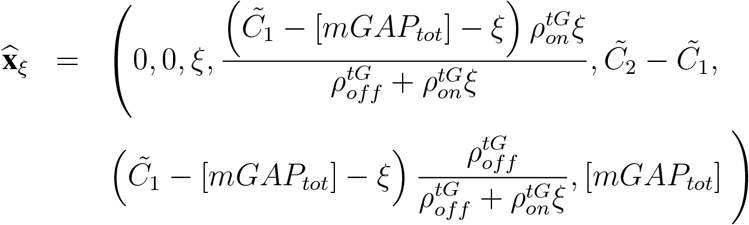

Case 2: 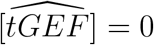 and 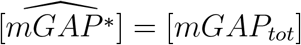

Using Eq. 3.39 to write 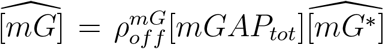 and subtracting Eq. 3.38 from Eq. 3.37, we obtain the expressions for [*mG*^∗^] and [*mG*]

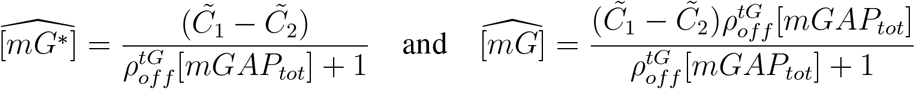

and thus we must have 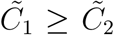. Now looking at Eq. 3.38 and substituting 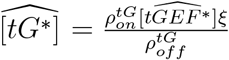 from Eq. 3.40, we obtain

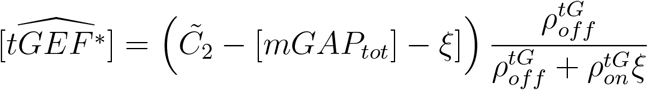

only if 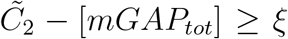. Therefore, in this case the *ξ*-dependent family of steady states is given by

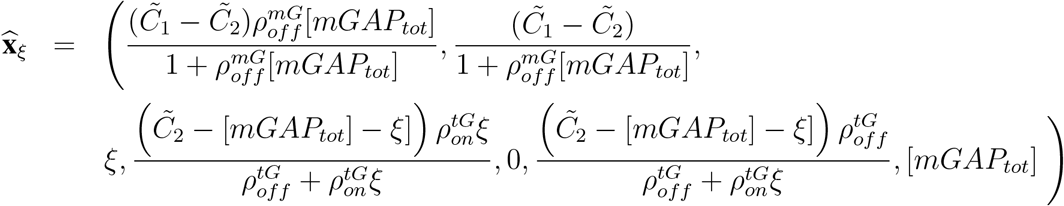

Case 3: 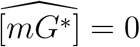 and 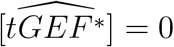

From Eqs. 3.39 and 3.40, we have 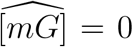 and 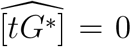, respectively. Subtracting Eq. 3.38 from Eq. 3.37, in this case we get 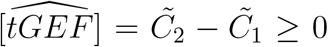 only if 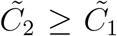. Now, from the conservation law given by Eq. 3.37, we obtain 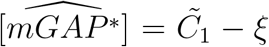 and thus 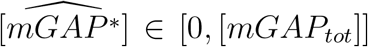 only if 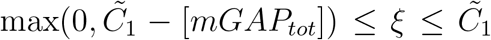. In this case, the *ξ*-dependent family of steady states is given by

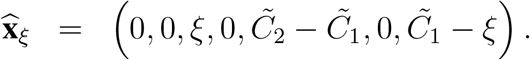

Case 4: 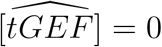 and 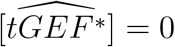

Eq. 3.40 gives 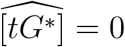 and the conservation law given by Eq. 3.38 yields 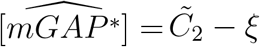. Now using Eq. 3.39 to write 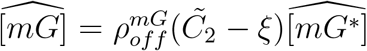, the conservation law given by Eq. 3.38 gives

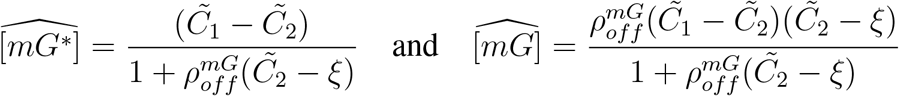

and since [*mGAP* ^∗^] ∈ [0, [*mGAP*_*tot*_]] and the steady states must be nonnegative, we must have

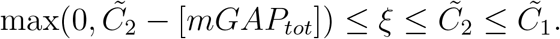

The *ξ*-dependent familiy of steady states is therefore given by

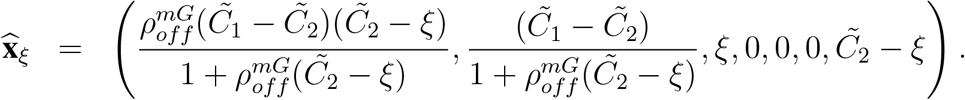

